# EMBER multi-dimensional spectral microscopy enables quantitative determination of disease- and cell-specific amyloid strains

**DOI:** 10.1101/2023.02.01.526692

**Authors:** Hyunjun Yang, Peng Yuan, Yibing Wu, Marie Shi, Christoffer D. Caro, Atsushi Tengeiji, Shigeo Yamanoi, Masahiro Inoue, William F. DeGrado, Carlo Condello

**Affiliations:** Institute for Neurodegenerative Diseases, University of California, San Francisco, CA 94143; Department of Pharmaceutical Chemistry, Cardiovascular Research Institute, University of California, San Francisco, CA 94158; Daiichi Sankyo Co. Ltd, Tokyo, Japan; Department of Neurology, University of California, San Francisco, CA 94143

## Abstract

In neurodegenerative diseases proteins fold into amyloid structures with distinct conformations (strains) that are characteristic of different diseases. However, there is a need to rapidly identify amyloid conformations *in situ*. Here we use machine learning on the full information available in fluorescent excitation/emission spectra of amyloid binding dyes to identify six distinct different conformational strains *in vitro*, as well as Aβ deposits in different transgenic mouse models. Our EMBER (excitation multiplexed bright emission recording) imaging method rapidly identifies conformational differences in Aβ and tau deposits from Down syndrome, sporadic and familial Alzheimer’s disease human brain slices. EMBER has *in situ* identified distinct conformational strains of tau inclusions in astrocytes, oligodendrocytes, and neurons from Pick’s disease. In future studies, EMBER should enable high-throughput measurements of the fidelity of strain transmission in cellular and animal neurodegenerative diseases models, time course of amyloid strain propagation, and identification of pathogenic versus benign strains.

**Significance:** In neurodegenerative diseases proteins fold into amyloid structures with distinct conformations (strains) that are characteristic of different diseases. There is a need to rapidly identify these amyloid conformations *in situ*. Here we use machine learning on the full information available in fluorescent excitation/emission spectra of amyloid binding dyes to identify six distinct different conformational strains *in vitro*, as well as Aβ deposits in different transgenic mouse models. Our imaging method rapidly identifies conformational differences in Aβ and tau deposits from Down syndrome, sporadic and familial Alzheimer’s disease human brain slices. We also identified distinct conformational strains of tau inclusions in astrocytes, oligodendrocytes, and neurons from Pick’s disease. These findings will facilitate the identification of pathogenic protein aggregates to guide research and treatment of protein misfolding diseases.

## Introduction

Amyloid fibrils are insoluble protein aggregates with a diverse range of biophysical properties, biological functions and association with human diseases. Their stability and resistance to degradation implicates them in: 1) adhesion and biofilm formation in bacteria^1^, 2) spore development in fungi^2^, 3) rubber biosynthesis in plants^3^, 4) chemical catalysis^4^, 5) materials^5^, and 6) systemic organ amyloidosis and neurogenerative diseases in humans^6^. Some of the most well-characterized amyloids are composed of amyloid-β (Aβ), tau or α-synuclein (α-Syn) proteins and associate with cell death and brain dysfunction in Alzheimer’s (AD) and Parkinson’s disease (PD), the most prevalent neurodegenerative diseases. These amyloidogenic proteins coalesce to form a cross β-sheet fibrils, which are observed as deposits in the brain.^7,8^ Each amyloidogenic protein is capable of adopting a number of different three-dimensional amyloid structures, each with distinct molecular repeating structures.^9,10,11^ Combined with biochemical and pathological processes such as post translational modifications (PTMs) or protease activity, these differences are known as conformational strains. ^12,13,14,15,16^ Like viral strains, different amyloid strains can propagate over multiple passages in animals or cell culture as in the classical prion mechanism.^17,18^ Also, different strains of disease-causing proteins such as Aβ and tau lead to different pathologies and localization in the brain. More generally, the biological, material, and chemical properties of amyloids depend critically on their conformations. Thus, there is a great need for a rapid method to differentiate distinct conformational strains of amyloids in brain tissues (derived from rodent models and human donors), cultured cells, and cell-free *in vitro* systems. The method developed here, which discriminates between amyloids with differing sequences and conformations, will benefit research on amyloids in any tissue type or biological system.

Structural methods to differentiate conformational strains are very labor intensive, generally involving biochemical or biophysical analysis (e.g., solids NMR^19,20,21^, protease susceptibility^22^, isotope-edited IR^23^, cryo-EM^24^, etc.). Moreover, these methods lose spatial biological context due to the stringent purification steps to extract amyloids. There are histological fluorescent dyes^25,26,27,28^ and clinical PET imaging probes^29,30,31^ retaining spatial information. Amyloid-staining dyes with spectral features that can be used as fingerprints to differentiate distinct conformational strains^32,33^, and cryo-EM structures^34,35,36,37^ have been highly successful in identifying the conformations of a number of disease-associated amyloids. Thus, while strain-sensing dyes do not provide direct structural information, these dyes are sensitive to conformational strain behavior and enable rapid *in situ* assessment. For example, oligothiophene dyes discriminate PrP strains^38^, Aβ strains in AD etiological subtypes and α-Syn strains in Parkinson’s disease and multiple system atrophy (MSA).^39,40^ However, they are difficult to synthesize, and their strain-sensing ability has not been quantified to find whether a single molecule can be used to distinguish wide range of amyloids. To address some of these limitations we previously developed a method, which utilizes multiple commercial dyes that individually have limited resolving power, but in aggregate were able to identify conformational strains when combined with principal component analysis (PCA).^41,42^ However, the collection of spectra from multiple dyes proved impractical, due to the need to either de-stain tissues between dye applications or to examine adjoining tissue slices, which limited spatial resolution.

Thus, we sought a technology that relies on only one single dye to sense a wide range of *in vitro* and biological amyloids. We show that the excitation/emission spectra from a single stain can provide a wealth of discriminating power when analyzed with advanced machine learning methods. The combination of excitation and emission spectra has only rarely been used to identify conformational strains, and there have been no attempts to automate the collection and analysis of data. Our EMBER (Excitation Multiplexed Bright Emission Recordings) workflow enabled high-throughput measurements of *in vitro* generated amyloids, allowing identification of amyloid strains within a set of six different amyloid types. It also enabled the analysis of brain slices from diseased tissues, showing for the first time, large *in situ* differences in the conformational strains of tau amyloids between multiple AD subtypes as well as cell-type specific and spatially resolved tau strains in Pick’s disease (PiD). This method should facilitate the measurement of the fidelity of transmission of conformational strains in cellular and neurodegenerative diseases animal models used in fundamental research and drug discovery. Moreover, this technology will allow measurement of time courses of aggregation and fibril formation in aqueous solution. Thus, EMBER has the potential to significantly increase the resolution and information content of any application of fluorescence imaging or microscopy in normal and pathogenic amyloids.

## Results

EMBER uses fluorescence microscopy to evaluate the excitation/emission (XM) spectra of amyloid-dye complexes, followed by PCA^43^ (principal component analysis), UMAP^44^ (uniform manifold approximation and projection) analysis, or Resnet-based neural network (NN)^45^ to identify and quantify spectral differences that are useful for differentiating amyloid strains (Fig 1). The method can be used on either tissue slices, cultured cells, or amyloid fibrils prepared *in vitro*. We began with *in vitro* fibrils, as they are relatively homogeneous, reproducible, and devoid of complicating cellular factors. In particular, we sought to discover dyes that can cleanly identify 6 different amyloids, Aβ40, Aβ42, α-Syn fibril, α-Syn ribbon, 0N3R tau and 0N4R tau. The fibrils were obtained by shaking the appropriate monomer from three to seven days to assure complete fibril formation. The process begins by mixing a dye with a suspension of a given amyloid strain in a microtiter well, and the plate is centrifuged to settle the amyloids to the bottom of the well. The concentration of fibrils is adjusted to provide a sparse collection of individual clumps of amyloid fibrils, which are imaged by a fluorescent microscope capable of measuring excitation and emission spectra. The spectra can be viewed as conventional overlay plots of emission intensity versus wavelength, each spectrum representing a different excitation wavelength. Alternatively, the individual emission spectra can be laid next to one another to create a sawtooth like profile (Fig. 1F), designated as the XM profile, which describes the full spectral details in a manner that allows easy visual comparison of dyes and subsequent analyses. XM profiles are measured for each particle in the well, and then this process is repeated for the six fibril types for a given dye. This results in approximately 300 XM profiles for a given fibril per micrograph, providing very rich structure sensitive set of data for probing amyloid strain behavior.

**Fig. 1.**
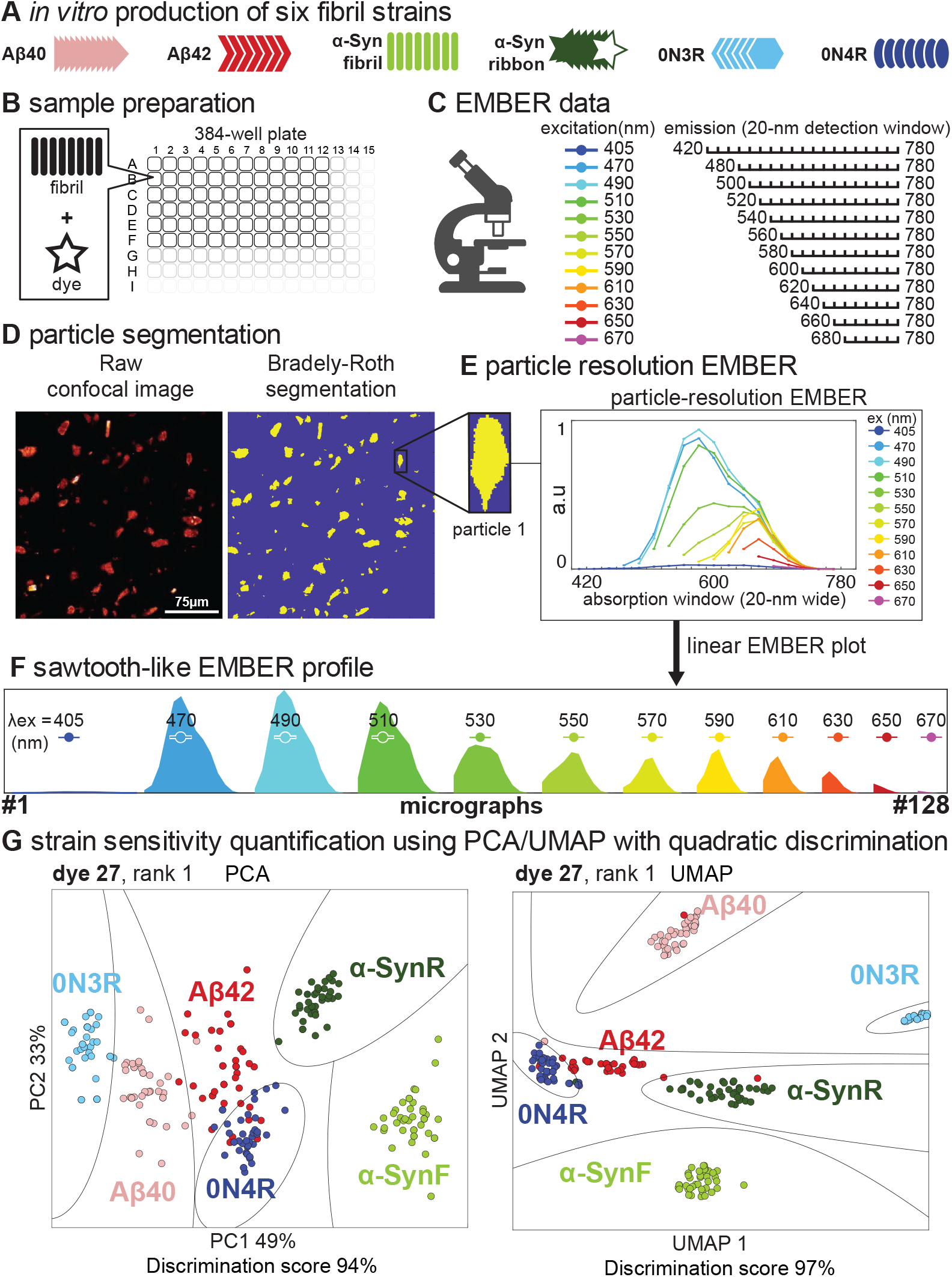
EMBER (Excitation Multiplexed Bright Emission Recordings) workflow. (A) Six distinct conformational fibrils are prepared. (B) Each of six *in vitro* prepared fibrils are mixed with each 145 dyes in a 384-well plate. (C) EMBERs of each fibril and dye mixture are collected. (D) Bradley-Roth segmentation is performed to provide a particle-resolution EMBER plot. (E & F) Overlay EMBER plot or linear sawtooth-like EMBER plot. (G) Individual EMBER plots are concatenated for PCA and UMAP analysis followed by quadratic discrimination to quantify conformational strain sensitivity of dye. Boundaries pertaining fit discriminants are presented in black lines.

We next use PCA, UMAP, and NN on the collection of XM profiles to determine how efficient the dye is at discriminating the six different fibril types in an unsupervised manner (Fig 2). The intensities of the XM profile are listed as a column vector, and a X*Y matrix is created in which X is the total number of particles across all 6 fibril types and Y is the number of intensities measured in a single XM profile. PCA and UMAP are then used to determine the variability of the XM profiles for each particle. The clustering of the points for a given type of fibril is useful in determining the degree of homogeneity of the sample. Points for additional fibril types that fall outside of a given fibrils cluster reflect differences in the environment of the bound dye. If the spectral features for a given fibril (e.g., Aβ42) are distinct from those of the other fibril types, the Aβ42 points will form an isolated cluster. We determine both PCA and UMAP (discrimination) plots. PCA is not as discriminating as UMAP but the Eigenvectors of PCA readily provide important physically meaningful information about which spectral features contribute most to discrimination. Thus, once a highly discriminating dye has been identified by the UMAP process, the Eigenvectors can inform the choice of excitation and emission wavelengths for more conventional imaging. Fig. 1 illustrates PCA and UMAP profiles for one particularly discriminating dye from our collection and Fig. S1 demonstrates strain-sensitivity and reproducibility of EMBER data collection over a three-day period by two operators.

**Fig. 2.**
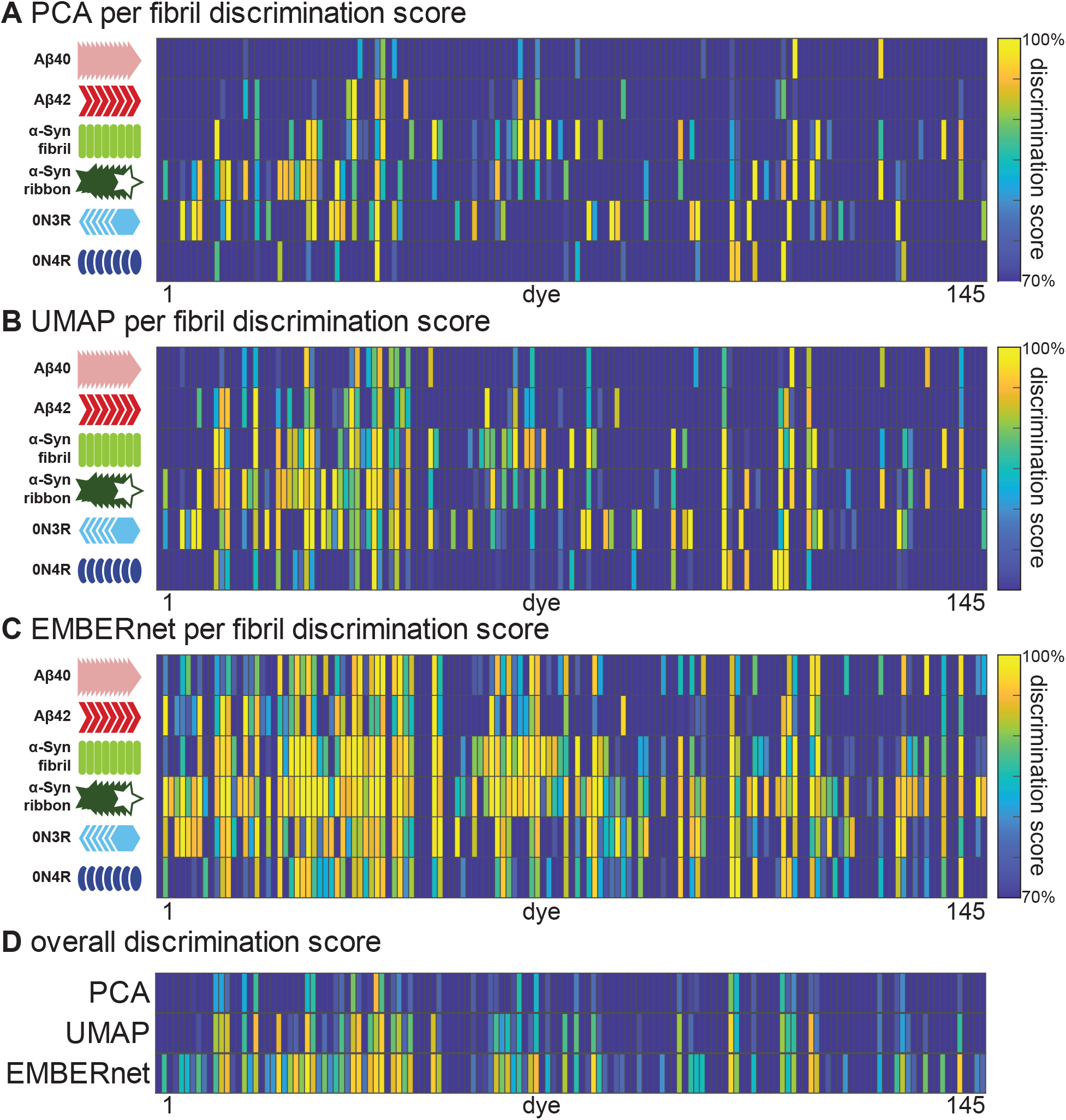
Per dye discrimination scores of each fibril type from (A) PCA (B) UMAP (C) EMBERnet and (D) across all fibril types. The heatmap plots are showing all discrimination scores >70% in the gradient colormap and all discrimination scores <70% in dark blue.

This process was then repeated for 145 different dyes, to find which individual dyes are most able to discriminate this collection of fibrils. The dyes comprise 56 loosely associated homologues of PBBs, 24 laser dyes, 14 curcumin homologues, 4 fsb homologues, 2 oligothiophene derivatives, and 47 other published or patented amyloid-binding dyes (Table S1). To aid the analysis, a MATLAB script was written to automate the segmentation and spectral measurement of individual particles. Using quadratic discrimination to determine the degree of overlap between clusters we identified 18 different dyes with discriminating power beyond 90% — a metric that ranges from 0% for no discrimination to 100% for full discrimination (Fig. 2, Fig. S2, and Table S1).

We also examined supervised methods of analysis. A convolutional neural net (EMBERnet) was trained on 80% of the XM profiles, which had been pre-assigned to each fibril type, retaining 10% each for validation and test sets (Fig 2 and Fig. S3–S5). As we expected, EMBERnet performed favorably relative to PCA and UMAP. For the most discriminating dyes, we observed very high discrimination scores using both EMBERnet and UMAP. However, EMBERnet performed better over a much wider range of dyes, indicating that it is better able to discover differences in even closely related spectra. Thus, EMBERnet is the method of choice when there is a large body of spectral data that has been assigned to each fibril type. On the other hand, as an unsupervised method UMAP shows its versatility in its ability to identify distinct clusters of conformers, even when the conformers have not been pre-assigned. This ability provides a powerful tool to discover systematic, unbiased differences between: 1) different amyloid preparations; 2) different cell types or 3) distinct spatially resolved inclusions within a single tissue slice.

### Discrimination of *in vitro* fibrils with EMBER

Fig. 3 illustrates typical spectra and discrimination plots for a low, intermediate, and highly resolving dyes. MCAAD-3 (dye 110) is one in the most discriminating group (98%); although its emission spectra look similar for the different fibrils, their intensity profiles do not vary uniformly with respect to excitation wavelength. Thus, a full EMBER analysis is able to identify dyes that do not have large spectral shifts, but nevertheless have very well-defined differences that are highly reproducible between fibrils of a given type – but vary reproducibly between different fibril types. Curcumin-stained XM exhibits quite different behavior, showing large differences in XM intensities between fibril types. The emission spectrum has two peaks whose relative intensities vary markedly with respect to the excitation wavelength in a fibril type-specific manner. Additionally, λ_ex max_ for the two peaks shift with the fibril type, providing an additional discriminating feature.

**Fig. 3.**
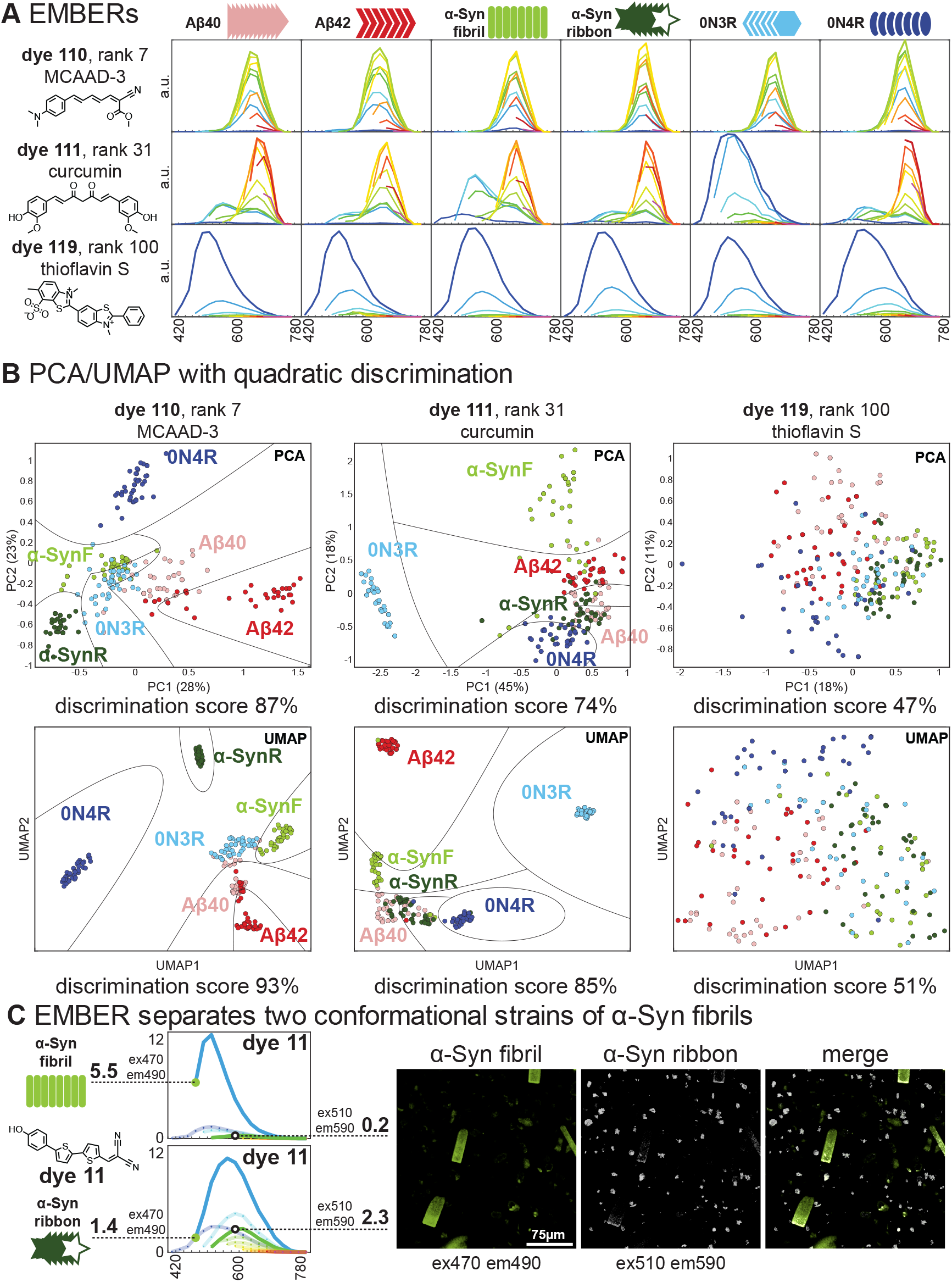
EMBERs of three dyes. (A) EMBERs of top, middle, low-tier dyes, and their chemical structures. (B) PCA and UMAP analysis of EMBERs. (C) Discrimination of α-SynF and α-SynR fibril with excitation and emission shift. α-SynF and α-SynR were mixed in a single well and dye 11 was added to achieve two different stained-fibrils in a single micrograph.

Such a dye might be ideal when a single excitation and/or emission is monitored, and it is not feasible to record full spectra. In contrast, low-discrimination dyes can be quite useful for broadspectrum staining. Indeed, the commonly used dye Thioflavin S was found to have a low discrimination score (51%, rank = 100th). Interestingly, in some cases, dyes perform reasonably well when excited at single wavelength, while the performance of other dyes only manifest in upon examination of both excitation/emission spectrum emphasizing the utility of the EMBER method (Fig. S6).

Thus, we used single dyes in our collection that could readily discriminate fibril types in a suspension, without the need for full spectral analysis (Fig. 3C). For example, we found that dye 11 stained α-Syn fibrils —α-SynF emitted brightly with an excitation/emission pair of 470/490 nm XM, while α-SynR was dim at these wavelengths, but quite bright with 530/590 nm XM. Satisfyingly, the two conformational strains of a single protein were easily identified using this method. Other pairs of XM selectivity can be readily extracted from Table S1 which outlines the XM_max_ pairs for given fibril type per dye.

### Quantification of conformational strain sensitivity in *ex vivo* mouse brain slices

We next asked if we could validate strain-sensing dyes discovered using our *in vitro* platform for recombinant fibrils in *ex vivo* applications. To evaluate if novel strain-sensing dyes could readily identify differences in *ex vivo* brain slices, we selected two different, well-characterized transgenic AD mouse models of human Aβ deposition: 1) Tg(APP23) mice which produce plaques rich in Aβ40 isotype and deposit slowly, and 2) Tg(5xFAD) mice which produce plaques rich in Aβ42 isotype and deposit rapidly. Using formalin-fixed mouse brain sections, we evaluated approximately 50 top scoring dyes (identified above) to assess strain sensitivity and tissue compatibility; 20 (of the 50) exhibited little non-specific background tissue staining and were selected for further examination. Qualitatively, the dyes could be separated into three classes: the first class stained a single strain and were bright over a range of excitation and emission spectrum. Dye 111 is typical of this class, as it brightly stains 5xFAD plaques with λ_ex_ = 405 nm and 470 nm but failed to appreciably fluoresce when it is used to stain APP23 plaques (Fig. 4A). *A* second class of dyes-stained plaques in both brain types, but the excitation wavelength required for detection was different between the two plaques. For example, dye 40 excites both plaque types with λ_ex_ = 405 nm but shows strong fluorescence for only APP23 when excited at 470 nm. Finally, some dyes, such as dye 27, stained plaques in a strain-independent manner over a range of wavelengths. Thus, our collection of dyes should be helpful, over a wide range of applications.

**Fig. 4.**
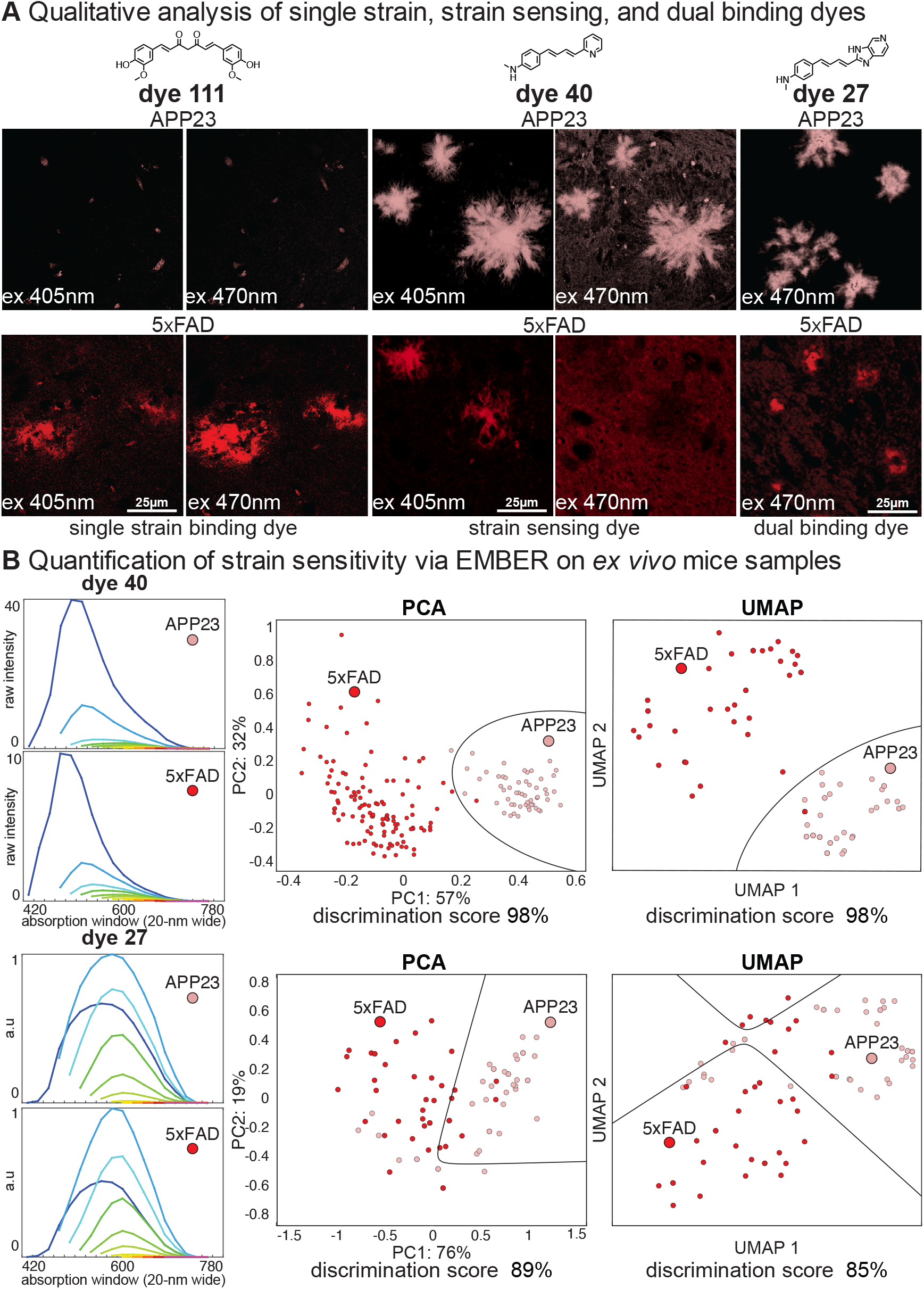
(A) Qualitative analysis of strain sensitivity against Aβ plaques formed in APP23 and 5xFAD tg AD mice models. (B) Dye strain sensitivity quantified via EMBER on *ex vivo* mice samples.

Further analysis of these data using the full EMBER pipeline showed additional resolving power. The plaques were identified, and spectra were measured using the same workflow as for the *in vitro* fibrils. In Fig. 4B, each particle on the PCA and UMAP plot represents one XM profile from a single plaque. The qualitative discrimination seen in Fig. 4A is consistent with the quantitative ability to discriminate (Fig. S7). Dye 40, which showed a higher fluorescence intensity for APP23 plaques over 5xFAD plaques at 470 nm excitation, had a 98% strain sensing score (based on UMAP) for the two brain types. Importantly, EMBER analysis was also able to provide some discrimination, even when the spectra appeared similar by visual examination as seen with dye 27 (Fig. 4B). This shows the power of EMBER and a library of dyes to solve the joint optimization problem of strain-sensing and tissue compatibility.

### Discrimination of Aβ plaques and neurofibrillary tau tangles in sAD brain samples

Using the top strain sensing dyes that were compatible with mouse tissue, we next identified dyes that can stain and differentiate both Aβ plaques and the tau neurofibrillary tangles (NFTs) in AD brain samples from patient donors (Fig. 5). Since these two deposit types are both present in a single AD brain slice, strain discrimination analysis under the same micrograph was possible. The plaques and tangles were initially assigned by visual analysis of their morphologies and were confirmed by antibody staining (Fig. S8). We focused on filamentous plaques as defined previously^46^, which were highly abundant relative to compact dense-core plaques, neuritic plaques, and cerebral amyloid angiopathy (CAA), and found consistently in all AD and DS brain samples we examined. Contrary to diffuse plaques, filamentous plaques bind can be labelled by amyloid-binding dyes. Consistent with previous findings with bf-188^32^, excitation at 405 nm resulted in high fluorescence intensity for Aβ and very low intensity at wavelengths greater than 561 nm. Just the opposite was observed for tau tangles. Thus, these dyes are ideally suited for identifying tau versus Aβ plaques with a single reagent in a single field of view, obviating the need for additional antibody staining.

**Fig. 5.**
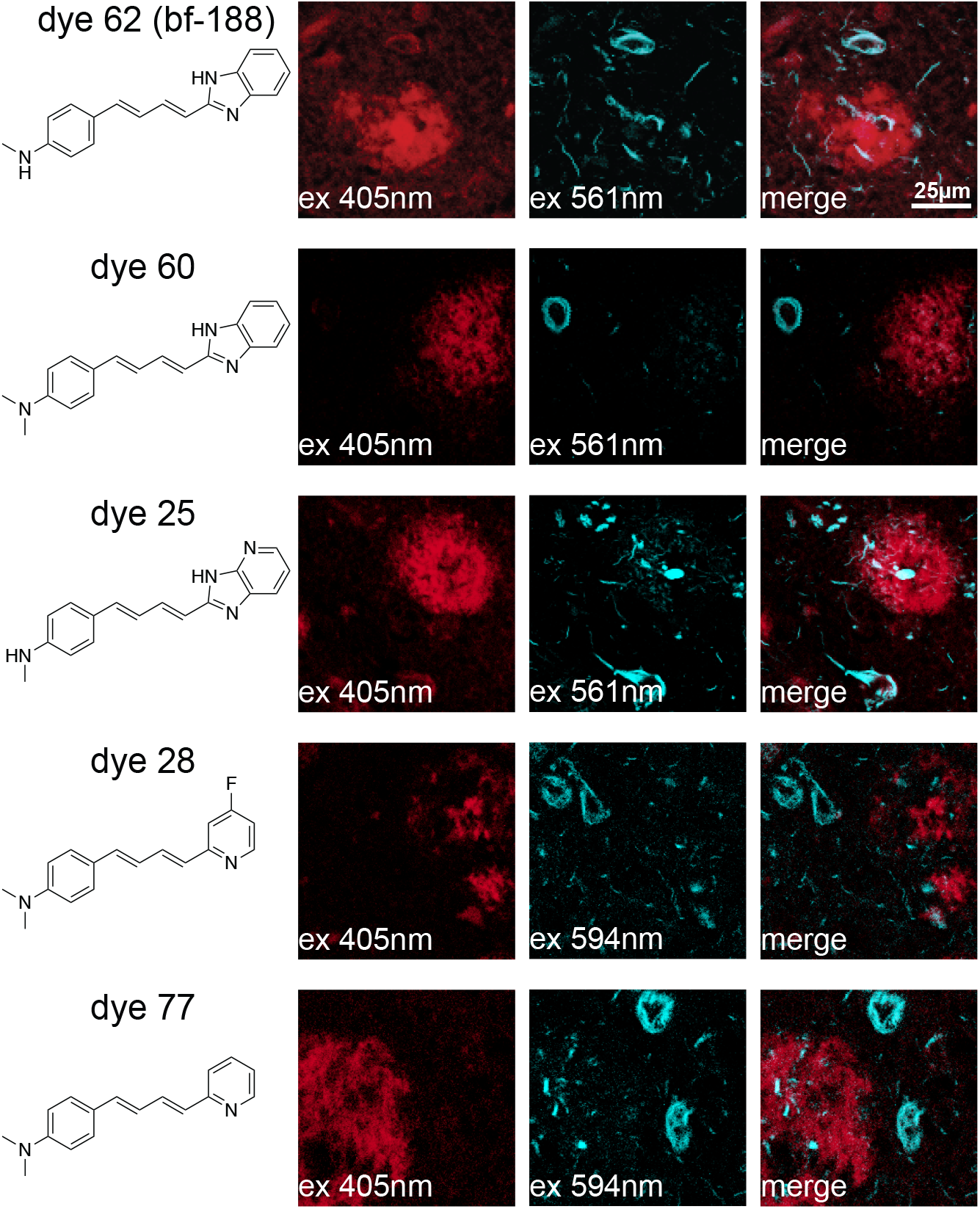
Structures of dyes that stain and strain sense Aβ plaque and tau tangle on sAD brain samples. Aβ plaques are excited at 405 nm excitation and tau tangles are excited at 561 nm excitation or above.

At the qualitative level, we looked for other dyes that can bind and discriminate between both deposit types by shifts in excitation. Of the top scoring dyes, we identified five dyes that spectrally separate the Aβ plaques and tau tangles. Dye 60, for example, favored Aβ plaque emission at 405 nm excitation compared to tau tangle emission at a more red-shifted excitation at 561 nm. Interestingly, all five dyes share chemical homology to PBBs and share the fluorescence properties of red-shifted XM for tau tangle and blue-shifted XM for Aβ plaque. We postulate that the shallow groove observed in the cryo-EM structures of tau fibril encourages dye interaction, causing the observed exciton coupling.^47,48,49^ Dye 60 was then selected for additional full EMBER analyses of Aβ plaques and tau tangles in a variety of different disease types.

### Aβ strain discrimination in Alzheimer’s disease and Down syndrome

We found dye 60 was particularly useful for discriminating conformational strains of Aβ plaques across four neurodegenerative diseases — sAD, fAD PSEN1, fAD APP, and Down syndrome (DS). Multiple brain samples from each of the neurodegenerative diseases (Table S2) were stained with dye 60 and their EMBER data was acquired from Aβ plaques (Fig. S9). Remarkably, plaques from each disease type form tight clusters in the plots (Fig. 6 and Fig. S10). Generally, we observed little inter-patient heterogeneity within a cohort even across fAD PSEN1 samples from three different brain banks for fAD PSEN1 (Fig. S11). The use of multiple excitation wavelengths increased the resolution, relative to single-wavelength measurements (Fig. S12).

**Fig. 6.**
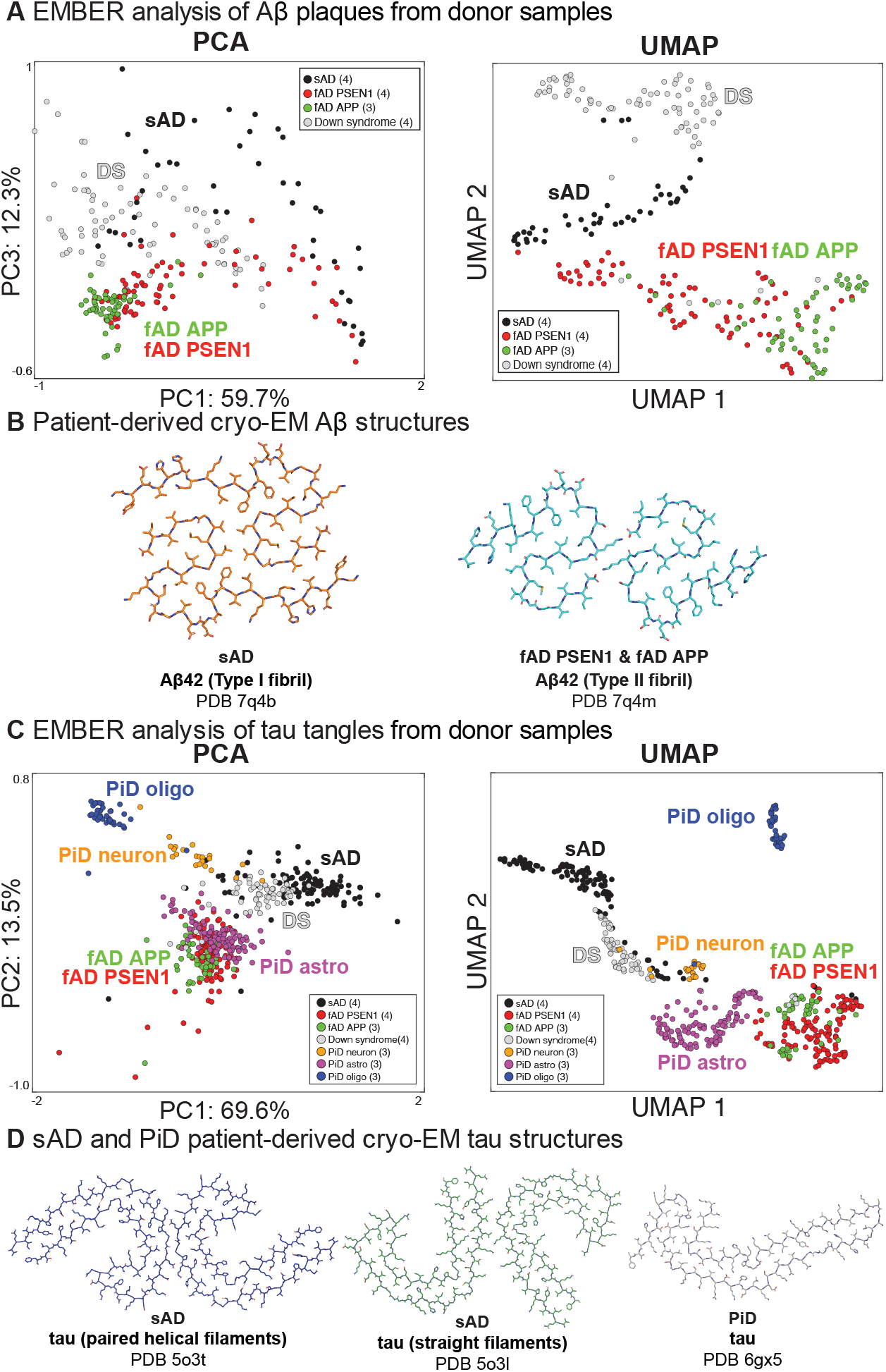
EMBER discriminates the conformational strains of Aβ plaques and tau tangles across neurogenerative diseases. (A) PCA and UMAP analysis of EMBERs of dye 60-stained Aβ plaques reveal inter-disease type cluster separation. fAD PSEN1 (red) and APP (green) share cluster in both PCA and UMAP. sAD (black) and Down syndrome (blue) share a cluster in PCA but are separated in the UMAP. Each particle represents one segmented Aβ plaque from EMBER micrograph. The number of donor samples in each cohort is within the parenthesis. (B) Cryo-EM structures of Aβ42 fibrils.^10^ (C) PCA and UMAP analysis of EMBERs of dye 60-stained tau tangles reveal inter-disease type cluster separation. fAD PSEN1 (red) and APP (green) share cluster in both PCA and UMAP. sAD (black), Down syndrome (gray), PiD neuron (orange), PiD astrocyte (purple), and PiD oligo (blue) form their own clusters depicting conformational strain polymorphism of tau tangles. Each particle represents one segmented tau tangle from EMBER micrograph. The number of patient samples in each cohort is within the parenthesis. (D) Patient-derived cryo-EM structures of tau fibrils.^9,34^

fAD PSEN1 involves a mutation in the gene encoding one of the proteolytic processing enzymes (γ-secretase) of the Aβ precursor protein (APP), and it results in a higher Aβ42/Aβ40 ratio than wildtype (WT). fAD APP is associated with a mutation to regions of APP that are outside of the Aβ peptide sequence, but within the γ-secretase cutting site. It too results in a higher Aβ42/Aβ40 ratio than WT. The structures of these two types of fAD had similar cryo-EM structures, which differed markedly from that of sAD.^10^ Thus, it is interesting that there is essentially no overlap (99% discrimination) between sAD (which has WT APP) and the two types of fAD. By contrast there is extensive overlap between fAD PSEN1 and fAD APP in the UMAP plots. Nevertheless, the fAD cohorts do show some separation, which might arise from conformational differences in the bulk of the sample, differences in the Aβ42/40 ratio or other associated molecules that are not apparent in highly purified samples used for cryo-EM.^50^

More impressively, the DS plaques were also fully separated from sAD and fAD plaques. Furthermore, we observed patient-specific (intra-DS group) clusters (Fig. S11), which is consistent with our prior work.^42^ High-resolution structures of Aβ fibrils from DS donors have yet to be identified; we postulate that the strain is similar but distinct to sAD as DS and sAD clusters overlap in the PCA and DS forms its own cluster in the UMAP. The PCA cluster area was larger for sAD and DS suggesting greater heterogeneity across the brain and amyloid deposits than that of the fAD samples.

### Quantitative determination of distinct tau strains within and across neurodegenerative diseases

Since dye 60 stains both Aβ plaques and neurofibrillary tau tangles (NFT), EMBER data from the tau tangles were collected and analyzed in the same micrographs (Fig. S13). In addition to the four disease types we analyzed for Aβ plaques (Table S2), we also examined tau deposits from the “tau-only” Pick’s disease (PiD) — cryo-EM structures of PiD tau fibrils are distinct from sAD tau fibrils.^34^ Moreover, tau deposits are found in three types of cells including neurons, astrocytes, and oligodendrocytes in PiD^51,52^, providing an opportunity to evaluate differences in conformational strains associated with different cell types.

As was the case for Aβ plaques, the tau tangles in DS and sAD were partially overlapping (Fig. 6). On the UMAP plot, five clusters are formed capturing the polymorphic nature of tau tangle strains across neurodegenerative diseases. sAD, DS, and fAD all form their own clusters. Interestingly, the familial mutations in APP and PSEN1 both lead to overproduction of Aβ42 in fAD, and the consequent tau tangles localize to the same cluster.

A clear difference was seen between the fAD and sAD cases. This finding contrasts with a cryo-EM study that suggests similar conformations for tau from fAD and sAD. It is not yet clear whether the difference we observe is a result of subtle differences in structure, as the cryo-EM structure of tau from fAD APP was at low resolution. Alternatively, the difference we see might arise from differences in PTMs or proteolysis. Interestingly, while we observed clear inter-patient variability of Aβ from fAD cases the tau tangles showed no interpatient variability (Fig. S14). Thus, different strains of Aβ plaques are capable of inducing the same strain of tau tangles. There was partial overlap between the UMAP clusters for DS and sAD, which is unsurprising given that aged DS patients often have co-morbid AD-like symptoms and the accumulation of tau tangles.^53^ This finding suggests that while there are large differences in the EMBER-detected conformational strains for Aβ plaques in DS vs. sAD, they ultimately lead to similar conformational strains of the tau tangles. Thus, the major difference in tau tangles appears to be temporal rather than structural.^54^

In addition to separating PiD tau deposits from all other neurodegenerative diseases tau in our analysis, EMBER was strikingly useful for discriminating between tau deposits localized to neurons, astrocytes, and oligodendrocytes in PiD brain samples. We confirmed the localization of tau deposits to each cell type using histochemical markers (Fig. S15) and their EMBER reproducibility (Fig. S16). These findings emphasize the power of EMBER to discriminate distinct strains and their spatial distribution in situ in multiple cell types (Fig. S17).

## Discussion

EMBER provides an attractive, well-automated, and objective approach for selection of strain-sensitive fluorescent dyes, and to maximize the strain information obtainable once a dye has been selected. The method is useful for examining very small samples with retention of spatial information. Although we have focused here on Aβ, tau, and α-Syn, the approach should be generalizable to address questions related to many other beneficial and pathologic biological amyloids^55^, as well as synthetic materials^56^. While we screened 145 dyes, the method can be easily extended to examine a larger library of commercial and custom dyes. It also provides a high degree of discrimination for selection of potential imaging agents that may differentiate only a single conformational form of the same protein. EMBER provides a very rapid method to probe the homogeneity of a sample, and when conformational information is available for a fibril type, it allows one to define a spectral signature for the structure. The method should also be highly useful for identifying dyes that are sensitive to conformation for *in vitro* or cell-free solution studies of intermediates and products for continuous monitoring of protein aggregation and amyloid formation on a fluorescent plate reader.

EMBER should also provide a highly discriminating probe for investigation of conformational fidelity during amplification or passage through cells and animal models. This microscopic approach provides a rapid conformational assessment, which can be correlated with cellular location of inclusions or their anatomical location. This will be particularly important for mechanistic studies and drug development, where it is important to confirm strain fidelity between multiple passages and time points.

While a fluorescent signature does not give direct structural information, it reports on the fine details of the binding site where the dye is located. As such, it not only reflects differences in conformation, but also might be influenced by PTMs, proteolysis, small molecule cofactors, and protein isoforms generated by differential gene splicing. EMBER is likely sensitive to each of these variables, and hence not entirely a measure of amyloid fibril structure, but also of these other features, which collectively define a conformational strain. For example, cryo-EM structures for sAD and fAD tau appear similar^57^, but different PTMs or truncated tau species not recognized by cryo-EM may contribute to the differences we observed in our EMBER data.

Interestingly, the spatial resolution afforded by EMBER revealed different conformational strains associated with tau deposits in different neural cell types in PiD samples. This finding shows that neurons, astrocytes, and oligodendrocytes have distinct conformational strains in the same brain. At first, our data may seem difficult to reconcile with the single PiD tau structure^34^ currently available from brain homogenates. However, this structure is derived from a single PiD donor and may only represent the predominate tau filament from the most abundant tau-laden cell-type present in the sample used for fibril purification. Consistent with observation, Falcon et al. have described a panel of western blots of purified tau extracts stained with different tau antibodies from nine PiD donors (including the one used for cryo-EM) that show varying band patterns and intensity differences (Fig. S6 from ref34). Thus, it will be interesting to determine whether the variation we see with cell type reflect differences in the conformation of the ordered amyloid core, or differences in PTM and the proteolytic variability seen previously.

We observed a dramatic increase in resolving power when we consider three spectral features that include excitation/emission spectra and overall brightness. The structural and chemical environment of the binding site affects the fluorescence according to a number of effects, including rigidity, excited state/ground state pKa, dye-fibril interactions, and orbital overlap between dyes held in close proximity along the regularly repeating structure of the amyloid. For example, in some cases, we observe a doublet in the emission spectra, which likely reflects excitonic coupling between stacks of aromatic dyes.^58^

Because EMBER relies on machine learning, it should be possible to expand EMBER to consider a number of different additional variables, each providing unique information. For example, dyes associate with differing amyloids at a range of affinities, so concentration should provide an additional discriminating variable. Moreover, the fluorescence lifetime and time-resolved Stoke’s shifts can be readily determined using instruments equipped with pulsed lasers and time-resolved spectral acquisition. Finally, the degree of immobilization of a dye within its binding site can be determined using steady-state and time-resolved fluorescence polarization.

In conclusion, EMBER is a highly resolving, automated method to discover and maximize the information of strain-sensing dyes with retention of spatial information. EMBER offers a facile quantitative approach that complements existing dye-based imaging methods to study conformational strains in cultured cells, animal models and primary human tissues. The method should be applicable to a variety of amyloids. In principle, it may also be useful for examine dyes that are responsive to molten globule-like intermediates and oligomers.^59^ Moreover, EMBER may be useful to rapidly screen hundreds of neurodegenerative disease donor samples and prioritize new targets for cryo-EM structural characterization.

## Materials and Methods

### Production of Aβ40 and Aβ42 fibrils

Aβ fibrils were produced following published protocols to produce homogenous preparations for structural determination.^60^ HPLC-purified and lyophilized Aβ40 and Aβ42 as TFA salts were purchased from rPeptide. 0.2 mg of lyophilized Aβ40 or Aβ42 was dissolved in 40 μL HFIP (5 mg/mL). The HFIP was evaporated overnight in the fume hood covered by Kimwipe and the resulting Aβ samples were speedvac for 30 min. The Aβ film was resuspended in 20 μL DMSO (10 mg/mL). The solution was briefly vortexed, water bath sonicated for 5 min, then diluted with 10 mM NaPhos buffer to 0.2 mg/mL final protein solution. The solution was pipette shaken in Thermomixer at 900 RPM for 72 h at 37 °C.

### Production of α-Syn fibril and ribbon

α-Syn fibrils were produced following published protocols to produce homogenous preparations.^61,62,63^ pET28a (kanamycin selected) vector encoding human α-synuclein was transformed into the E. coli strain BL21(DE3). Bacteria were grown to OD600=0.8 then IPTG induced for 3 h. The cells were pelleted, resuspended in 100 mL osmotic shock buffer (20 mM Tris-HCl, 40% sucrose, 2 mM EDTA, pH 7.2), incubated at room temp for 10 min, then centrifuged at 12k rpm for 20 min. The supernatant was decanted then the pellet was resuspended in 100 mL cold water containing 40 μL saturated MgCl2. The solution was left on ice for 3 min and centrifuged at 12k rpm for 20 min. The resulting supernatant was collected and lyophilized. The lyophilized powder was dissolved in 20 mL 5% acetonitrile (MeCN) containing 0.1% TFA, then filtered through 0.45-μm polypropylene syringe filter and purified on a semi-prep C4 HPLC column (H2O/MeCN containing 0.1% TFA; 5% to 95% MeCN over 30 min). The pure α-synuclein fractions were collected and lyophilized. To prepare α-Syn fibrils, lyophilized α-synuclein was dissolved in 50 mM Tris-HCl, pH 7.5, 150 mM KCl to result in a final concentration of 100 μM α-Syn. To prepare α-Syn ribbons, lyophilized α-synuclein was dissolved in 5 mM Tris-HCl pH 7.5 to result in a final concentration of 100 μM α-Syn. Both fibrillization conditions were shaken in a Thermomixer at 1000 rpm for 7 d at 37 °C. The fibril quality was controlled with YFP-labelled cell aggregation assay, sedimentation assay, and TEM imaging.

### Production of 0N3R tau and 0N4R tau fibrils

Tau fibrils were produced following published protocols to produce homogenous preparations for structural determination.^64,65^ The E. coli strain Rosetta (DE3) was transformed with pET28a (kanamycin selected) vector encoding 0N3R tau and 0N4R tau. The bacteria were grown to OD_600_=0.8 then IPTG induced for 6 h. Cells were pelleted and resuspended in 300 mL lysis buffer containing 20 mM MES (pH 6.8), 1 mM EGTA, 0.2 mM MgCl2, 5 mM DTT, and 1xcOmplete™ protease inhibitor cocktail (Roche). Cells were lysed with microfluidizer then boiled for 20 min, and the resulting solution was spun at 24,500 g. The supernatant was purified with a cation exchange column (self-packed with SP Sepharose Fast Flow resin, GE Healthcare) then further purified with a reverse-phase HPLC equipped with ZORBAX 300SB-C3 column (MeCN gradient from 5-50% over 45 min). HPLC fractions containing pure 0N3R tau and 0N4R tau were combined and lyophilized to result in a protein powder as TFA salts. The fibrillization of both 0N3R and 0N4R tau were performed in a 1.5-mL Eppendorf tube at 0.4 mg/mL tau concentration with 0.125 mg/mL heparin (8,000-25,000 Da). 0N4R tau was fibrillized in 1x PBS containing 1 mM DTT and 0N3R tau was fibrillized in 1x PBS containing 1 mM TCEP (pH 7). The solution was shaken at 37°C and 1400 rpm for 3 d. The fibril quality was controlled with aggregation assay, sedimentation assay, trypsin-digested SDS-PAGE gel, and TEM imaging.

### Plate-based *In vitro* fibril staining for high-throughput EMBER microscopy

Each dye was dissolved in 1x PBS buffer then centrifuged at 13,200 rpm to result either 25 μM or saturated dye solution. A 50-μL dye solution was mixed with 1-μL fibril solution which contained ~0.1 μg fibril in a 384-well plate (Corning™ BioCoat™ 384-Well, Collagen Type I-Treated, Flat-Bottom Microplate). Each well was mixed by pipetting up and down then the plate was centrifuged at 50 x g to form fibril pellets.

### EMBER microscopy data collection

The 384-well plate containing the stained fibrils were imaged with Leica SP8 confocal microscope using a 40x water immersion lens (1.1 NA), a white light and 405 nm lasers, a HyD detector at 512×512-pixel resolution at 0.75x zoom. For high-throughput data acquisition sequential data collection was achieved using the *Live Data* mode, a module in in the Leica LAS X software. For each field-of-view experiment, the optical plane was auto-focused with the highest sensitivity setting. To reduce the background noise from the bottom of the plate well, LightGate was set to 0.5–18 ns. First, a total of 110 images were acquired using in the Λ/λ-scan mode with excitations of 470, 490, 510, 530, 550, 570, 590, 610, 630, 650 and 670 nm wavelengths. The emission detection range started at 10 nm greater than a given the excitation wavelength, and ended at 780 nM, with 20-nm window. For example of 470 nm excitation, the images were collected at 480–500, 500–520, 520–540, 540–560, 560–580, 580–600, 600–620, 620–640, 640–660, 660–680, 680–700, 700–720, 720–740, 740–760, 760–780 nm. Then in the λ-scan mode, 18 additional images were collected at 405 nm excitation with emission detection intervals of 20-nm for 420 nm to 780 nm. For *ex vivo* brain sample data collection, the zoom was increased to 2.0 and FOVs were manually focused.

### Post-processing and particle segmentation

We developed a set of custom scripts in MATLAB to process the raw fluorescent images and segment the aggregated protein deposits. In brief, we applied locally adaptive thresholding strategy. First, we generated a projection from the EMBER dataset. The maximum intensity projection was calculated across the λ stack: for the XY coordinate of a pixel, the highest fluorescence intensity within the corresponding coordinate through the λ stack was selected, resulting in a 2D image. The Bradley-Roth image thresholding method was applied on the resulting image: the image was divided into approximately eight smaller neighborhoods, each with an independent thresholding value calculated from the local mean fluorescence intensity; each pixel was then assigned a binary background or foreground value based on its neighborhood threshold value. Foreground noise was reduced by applying an image erosion calculation with a disk-shaped structuring element. Background noise was reduced by applying a flood-fill algorithm to fill holes in objects. Parameters for the segmentation processes were determined with a trial dataset and kept constant for each experiment. All deposits were then overlaid on the raw images for inspection and incorrect segmentations were removed from downstream analysis.

### PCA and UMAP with quadratic discriminant classification

The signal-processing algorithm for the analysis of particle resolution EMBER spectra was executed in MATLAB with *pca*. Each identified EMBER particle from the particle segmentation was normalized to [0, 1] and then concatenated in an array for PCA. The principal component scores PC1 and PC2 were plotted. UMAP was performed in MATLAB with *run_umap* with default settings. The cluster classification algorithm for the analysis of PCA and UMAP plot was executed in MATLAB with *fitedise*. From the PCA or UMAP plot, 40 random particles from each fibril were concatenated in an array and grouped for quadratic discrimination. This process was repeated ten times and the average of ten accuracy scores was used as the discrimination score.

#### EMBERnet data preparation

This neural network problem was formulated as an image classification problem that is solved using EMBERnet, a ResNet-based deep learning architecture.^45^ An image was constructed for each particle from EMBER dataset creating a size 12 × 18 × 64 image representing 12 excitations and 18 emission windows. The image was resized to 12 × 128 × 128 with *OpenCV* with interpolation from the original set of pixels.^66^ The final images were fed for deep learning model.

#### EMBERnet model architecture

The basic unit of neural network architecture is a residual block, which consists of two convolutional layers^67^, a batch-normalization layer^68^, and a rectified linear unit (ReLU)^69^. Residual blocks were also applied to overcome the problem of exploding/vanishing gradients due to increasing the depth of neural networks.^45^ Finalized EMBERnet model consisted of 22 layers — 20 convolutional layers and 2 multi-layer perceptron (MLP) layers. The skip connection was used to develop deep network with 20 layers, and also increased the convergence speed during training.^45^

#### EMBERnet model training and testing

To find the optimum discrimination score of EMBER dataset, 10-fold Cross-Validation (CV) was utilized.^70^ The EMBER dataset was randomly shuffled and then split in to 10 folds (0–9-folds) to ensure the distributions of the six fibril types are similar. For the training, validation, and test set, we utilized 8-folds (0–7-folds), 1-fold (fold-8), and 1-fold (fold-9) respectively. The models were trained for the maximum 120 epochs with a batch size of 32 on one NVIDIA GPU. During the training, cross-entropy loss between the target and predicted outputs was minimized using Adam optimizer.^71^ The learning rate of the optimizer was set to 0.00001 at the beginning of the training, and then reduced by a factor of 10 after approximately 100 epochs if no improvement in the discrimination score was observed after 3 epochs. The model training is then terminated if no performance improvement was observed after 6 epochs. After the performance of the training, validation, and test sets were confirmed, we finalized the EMBERnet and moved onto EMBERnet validation by shifting the folds by 1. Here, we used the folds-1–8 for the training set, fold-9 as the validation set, and fold-0 as the test set, and the EMBERnet performance was similar validating our EMBERnet training and performance. The code for EMBERnet is available per request.

### Mouse brain samples

All comparisons were conducted on brains from age-matched animals. Animals were maintained in a facility accredited by the Association for Assessment and Accreditation of Laboratory Animal Care International in accordance with the Guide for the Care and Use of Laboratory Animals.^72^ All procedures for animal use were approved by the University of California, San Francisco’s Institutional Animal Care and Use Committee. Tg(APP23) mice, which express human APP (751-aa isoform) containing the Swedish mutation under the control of the Thy-1.2 promoter, were maintained on a C57BL/6 background.^73^ Tg(5xFAD) mice, which express human APP containing the Swedish, Florida, and London mutation with PSEN1 harboring M146L and L286V mutation under the control of the Thy-1.2 promoter, were maintained on a C57BL/6 background.^74^ In this study, Tg(APP23) mice were 24-months old and Tg(5xFAD) mice were 12-months-old. Mouse brain samples were harvested, immersion-fixed in 10% buffered formalin, and then embedded in paraffin following standard procedures.

### Dye staining in brain sections for EMBER microscopy

Formalin-fixed paraffin embedded (FFPE) mouse brains were sectioned (8-μm thickness) and glass mounted. To reduce the autofluorescence signals by greater than 90% intensity (e.g., lipofuscin or hemosiderin), FFPE slides were photobleached up to 48 h using a multispectral LED array in the cold room overnight to reduce the autofluorescence in the brain tissue.^75^ The sections were deparaffinized, PBS-washed and stained with 25 μM for 30 min. The sections were washed with PBS buffer and a coversliped with PermaFluor (Thermo) as the mounting media. For FFPE human brain sections, the same procedures were followed. For EMBER data collection, the same steps were taken without the autofocus function and with zoom of 1.5.

## Data availability

The data that supports the findings of this study including dye synthetic schemes are available from the corresponding author, upon reasonable request. The custom developed MATLAB code for micrograph particle segmentation and EMBER analysis is available here: (https://www.dropbox.com/sh/w5xtpciyy8u0loe/AACKPlHHIDkotAS3_xcq5oaPa?dl=0).

## ACKNOWLEDGEMENTS

This work was supported by the National Institutes of Health (AG061874 and AG002132). H.Y. received postdoctoral fellowship funding from the BrightFocus Foundation (A2020039F). Tissue samples were supplied by Professor William W. Seeley (UCSF Memory and Aging Center, San Francisco, CA) and the UCSF Neurodegenerative Disease Brain Bank, which is supported by NIH grants AG023501 and AG019724, the Tau Consortium, and the Bluefield Project to Cure FTD; Professor Elizabeth Head and the University of California Alzheimer’s Disease Research Center (UCI-ADRC, Irvine, CA), which is funded by NIH/NIA (P30AG066519); the London Neurodegenerative Diseases Brain Bank (King’s College London, England), which receives funding from the Medical Research Council UK and through the Brains for Dementia Research Project (jointly funded by the Alzheimer’s Society and Alzheimer’s Research UK); Dr. Andrew C. Robinson and the Manchester Brain Bank (University of Manchester, England), which is part of the Brains for Dementia Research program, jointly funded by Alzheimer’s Research UK and the Alzheimer’s Society; the Parkinson’s UK Brain Bank at Imperial College London; Professors Dennis W. Dickson and Michael DeTure and and were from the Brain Bank for Neurodegenerative Disorders at Mayo Clinic (Florida), which is supported by The Rainwater Charitable Foundation and the Mangurian Foundation; Professor Matthew P. Frosch and the Massachusetts Alzheimer’s Disease Research Center (director, M. P. Frosch), which received financial support from the NIH (P50AG005134). We received the CRANAD series of dyes as a gift from Professor Chongzhao Ran at MGH/Harvard University, and oligothiophene dyes as a gift from Professor K. Peter Nilsson at Linköping University (Sweden).

## Author contributions

H.Y., W.F.D and C.C. designed the research; H.Y., C.D.C., M.S., and C.C. performed the experiments; A.T., S.Y., and M.I. synthesized novel small-molecule dyes. H.Y and P.Y. designed and wrote custom MATLAB code; Y.W. designed and performed the convolutional neural net; H.Y., W.F.D., and C.C. analyzed the data; and H.Y., W.F.D. and C.C. wrote the paper. W.F.D. and C.C. supervised the study. All authors read and approved the manuscript.

