## supporting information for "EMBER multi-dimensional spectral microscopy enables quantitative determination of disease- and cell-specific amyloid strains"

### Table of Contents

**Fig. S1.** EMBER reproducibility study against *in vitro* fibrils. S1

**Fig. S2.** Randomization and quadratic discrimination of *in vitro* fibril data. S2

**Fig. S3.** Data preparation for EMBERnet. S3

**Fig. S4.** EMBERnet model architecture. S4

**Fig. S5.** EMBERnet model training and testing. S5

**Fig. S6.** EMBER vs single-wavelength excitation comparison against *in vitro* fibrils. S6

**Fig. S7.** Randomization and quadratic discrimination of plaque data in Tg mice. S7

**Fig. S8.** Validation of dye 60 dual XM properties in sAD sample using immunohistochemistry. S8

**Fig. S9.** EMBER plots of Aβ plaques across neurogenerative samples. S9

**Fig. S10.** PC1/PC2 and PC1/PC3 of Aβ plaque EMBERs across neurodegenerative diseases. S10

**Fig. S11.** Inter-patient heterogeneity plot for Aβ plaques. S11

**Fig. S12.** EMBER vs single-wavelength excitation comparison for Aβ plaques. S12

**Fig. S13.** EMBER plots of tau tangles across neurodegenerative samples. S13

**Fig. S14.** Inter-patient heterogeneity plot for tau tangles. S14

**Fig. S15.** Validation of dye 60 cell-type specific labeling in PiD sample using immunohistochemistry. S15

**Fig. S16.** EMBER reproducibility study against Pick astrocytes. S16

**Fig. S17.** EMBER vs single-wavelength excitation comparison for tau tangles. S17

**Table S1**. Dye structure, name, discrimination score, and $\lambda_{max}$ em/ex against *in vitro* fibrils.

**Table S2**. Sources of postmortem human brain tissue samples.

**Data S1.** *In vitro* fibril EMBER – raw data.

**Data S2.** *In vitro* fibril EMBER – normalized data.

**Data S3.** *In vitro* fibril PCA plots.

**Data S4.** *In vitro* fibril UMAP plots.

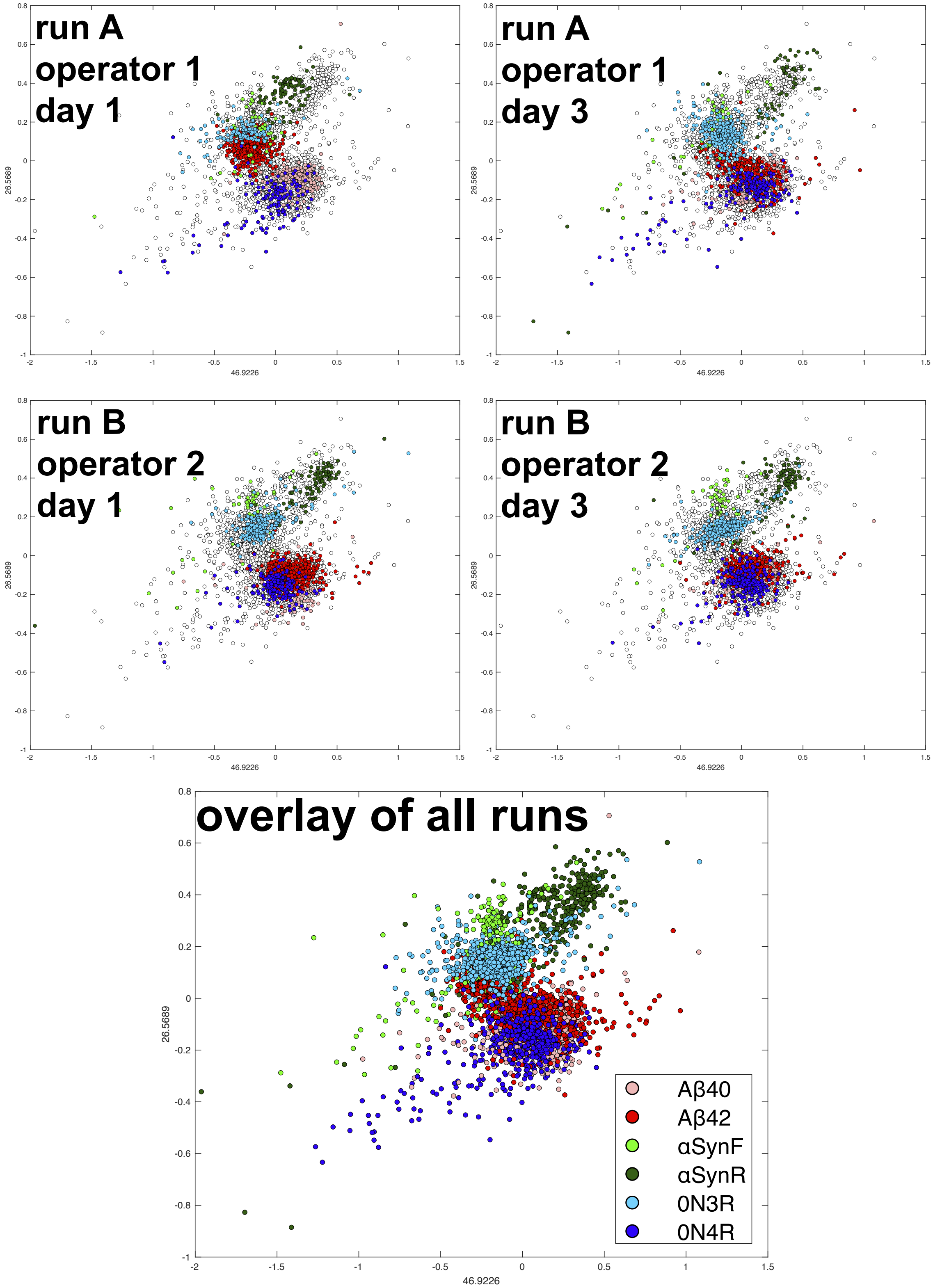


**Fig. S1.** EMBER reproducibility of six *in vitro* fibril types. Two operators (1 and 2) each prepared samples and collected EMBER dataset (run A and B) against six *in vitro* fibrils (Aβ40, Aβ42, αSynF, αSynR, 0N3R, 0N4R). EMBER data for each run was recollected at day 3. All four individual EMBER datasets were combined in an array and PCA was performed. PCA plots of overlay as well as the four individual runs are shown above.

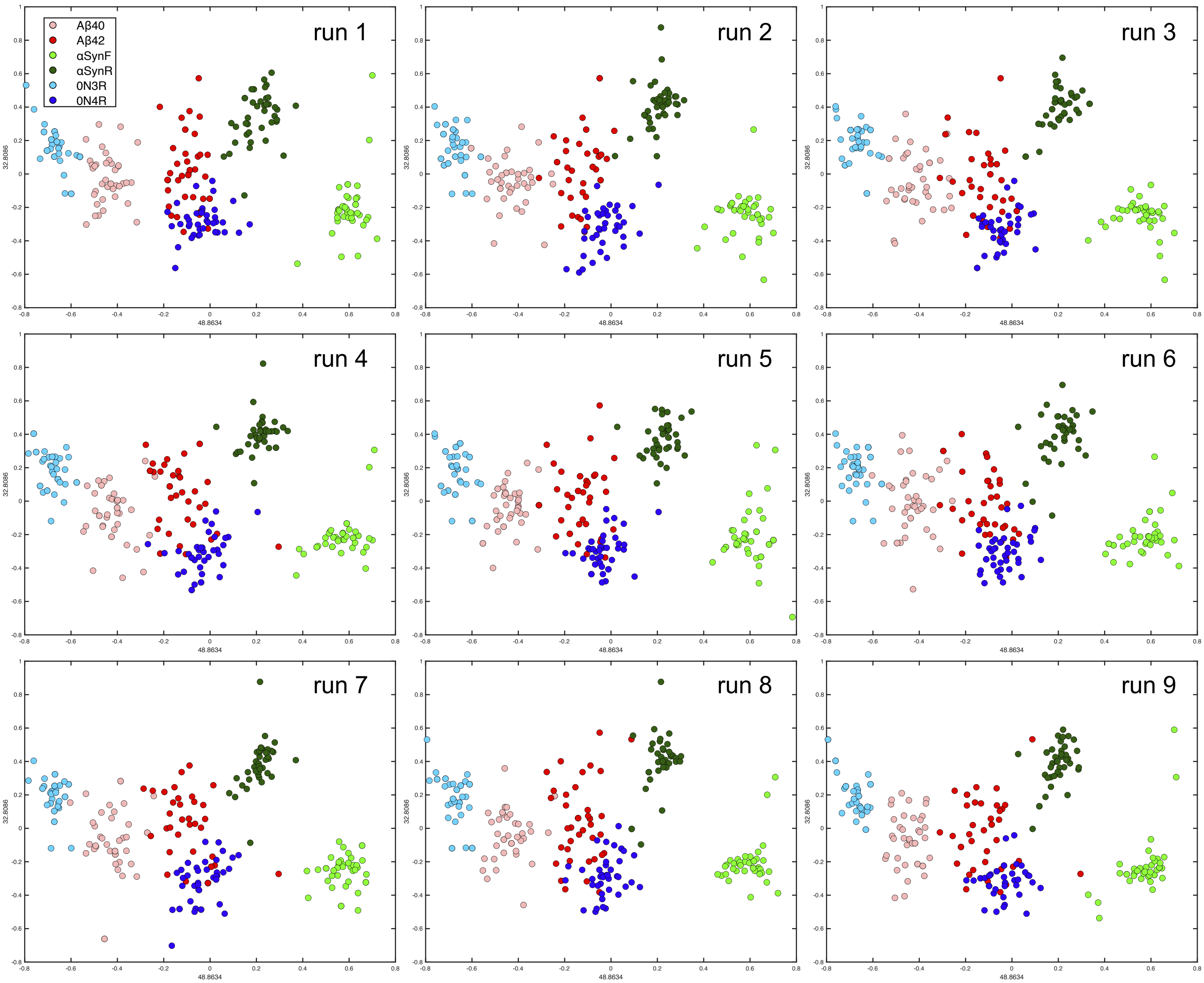


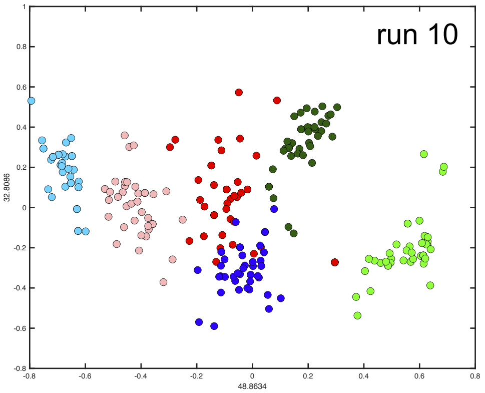


| Run # | Discrimination scores | Average discrimination scores |
|---|---|---|
| run 1 | 93% | 92% |
| run 2 | 91% | |
| run 3 | 94% | |
| run 4 | 93% | |
| run 5 | 94% | |
| run 6 | 92% | |
| run 7 | 93% | |
| run 8 | 92% | |
| run 9 | 91% | |
| run 10 | 90% | |

**Fig. S2.** 10 randomized data point selections and quadratic discrimination for six *in vitro* fibril types. Ten repetitions of quadratic discrimination cluster classification algorithm were performed to quantify discrimination score against *in vitro* fibrils. For PCA or UMAP plot, 40 random particles from each fibril dataset were concatenated into an array and grouped for quadratic discrimination. Discrimination score is calculated per run as shown above, and the average of ten discrimination scores was used as the discrimination score as presented in the main Figure 1.

(a)

| | excitation | 405 | 470 | 490 | 510 | 530 | 550 | 570 | 590 | 610 | 630 | 650 | 670 |
|---|---|---|---|---|---|---|---|---|---|---|---|---|---|
| emission | | 0 | 1 | 2 | 3 | 4 | 5 | 6 | 7 | 8 | 9 | 10 | 11 |
| 420 | 0 | 110 | | | | | | | | | | | |
| 440 | 1 | 111 | | | | | | | | | | | |
| 460 | 2 | 112 | | | | | | | | | | | |
| 480 | 3 | 113 | 0 | | | | | | | | | | |
| 500 | 4 | 114 | 1 | 15 | | | | | | | | | |
| 520 | 5 | 115 | 2 | 16 | 29 | | | | | | | | |
| 540 | 6 | 116 | 3 | 17 | 30 | 42 | | | | | | | |
| 560 | 7 | 117 | 4 | 18 | 31 | 41 | 54 | | | | | | |
| 580 | 8 | 118 | 5 | 19 | 32 | 40 | 55 | 65 | | | | | |
| 600 | 9 | 119 | 6 | 20 | 33 | 39 | 56 | 64 | 75 | | | | |
| 620 | 10 | 120 | 7 | 21 | 34 | 38 | 57 | 63 | 76 | 84 | | | |
| 640 | 11 | 121 | 8 | 22 | 35 | 37 | 58 | 62 | 77 | 85 | 92 | | |
| 660 | 12 | 122 | 9 | 23 | 36 | 36 | 59 | 61 | 78 | 86 | 93 | 99 | |
| 680 | 13 | 123 | 10 | 24 | 37 | 35 | 60 | 60 | 79 | 87 | 94 | 100 | 105 |
| 700 | 14 | 124 | 11 | 25 | 38 | 34 | 61 | 59 | 80 | 88 | 95 | 101 | 106 |
| 720 | 15 | 125 | 12 | 26 | 39 | 33 | 62 | 58 | 81 | 89 | 96 | 102 | 107 |
| 740 | 16 | 126 | 13 | 27 | 40 | 32 | 63 | 57 | 82 | 90 | 97 | 103 | 108 |
| 760 | 17 | 127 | 14 | 28 | 41 | 31 | 64 | 56 | 83 | 91 | 98 | 104 | 109 |

(b)

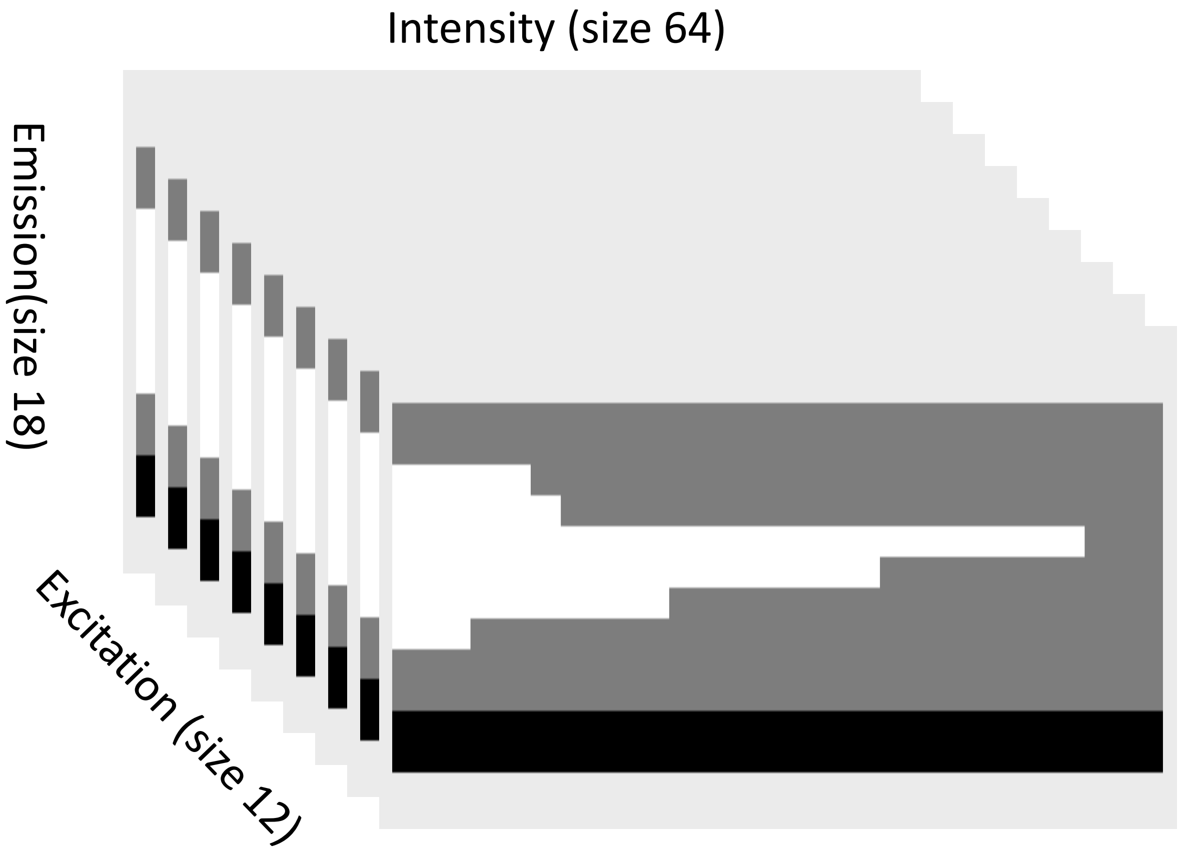


**Fig. S3.** The problem was formulated as an image classification problem that was investigated by an in-house ResNet-based deep learning architecture. (a) EMBER data format consisted of 12 excitations and 18 emission windows. (b) The constructed image for each experiment with size 12 x 18 x 64. The images were then resized to 12 X 128 X 128 by interpolating the pixel from the original set of pixels and were fed for the deep learning model.

(a) The basic residual unit

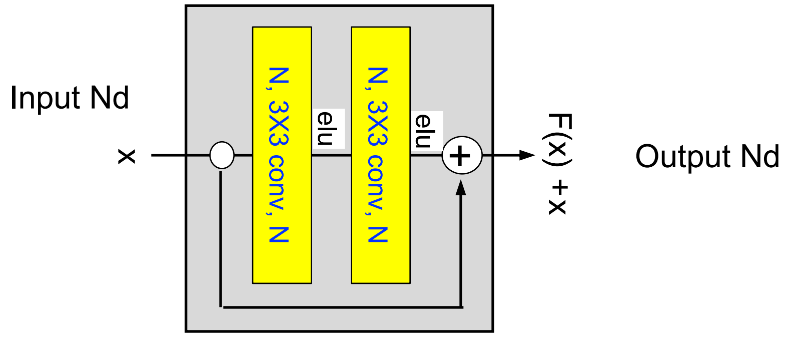


[A convnet layer:  layer # in channels, filter size, # out channels]

(b) Network Architecture

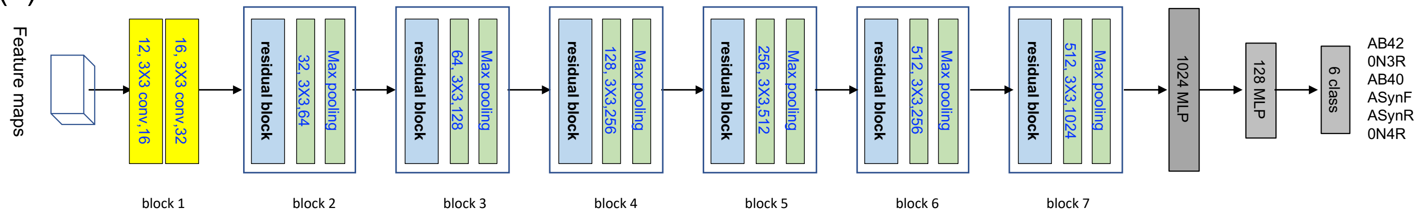


**Fig. S4.** Outline of ResNet-based 20-layer convolution neural network for multi-class image classification for each fibril types and all fibril types.

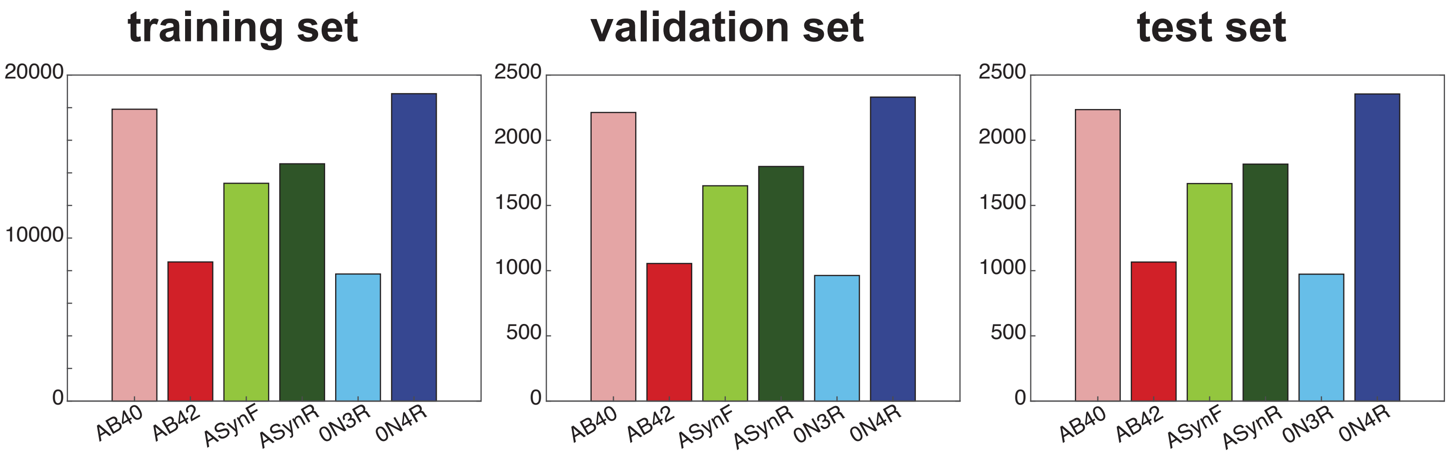


**Fig. S5.** The sample distribution of six fibril types in the first of 10 runs, where fold 0 to 7 are pooled as train set, fold 8 as validation set, and fold 9 as test set.

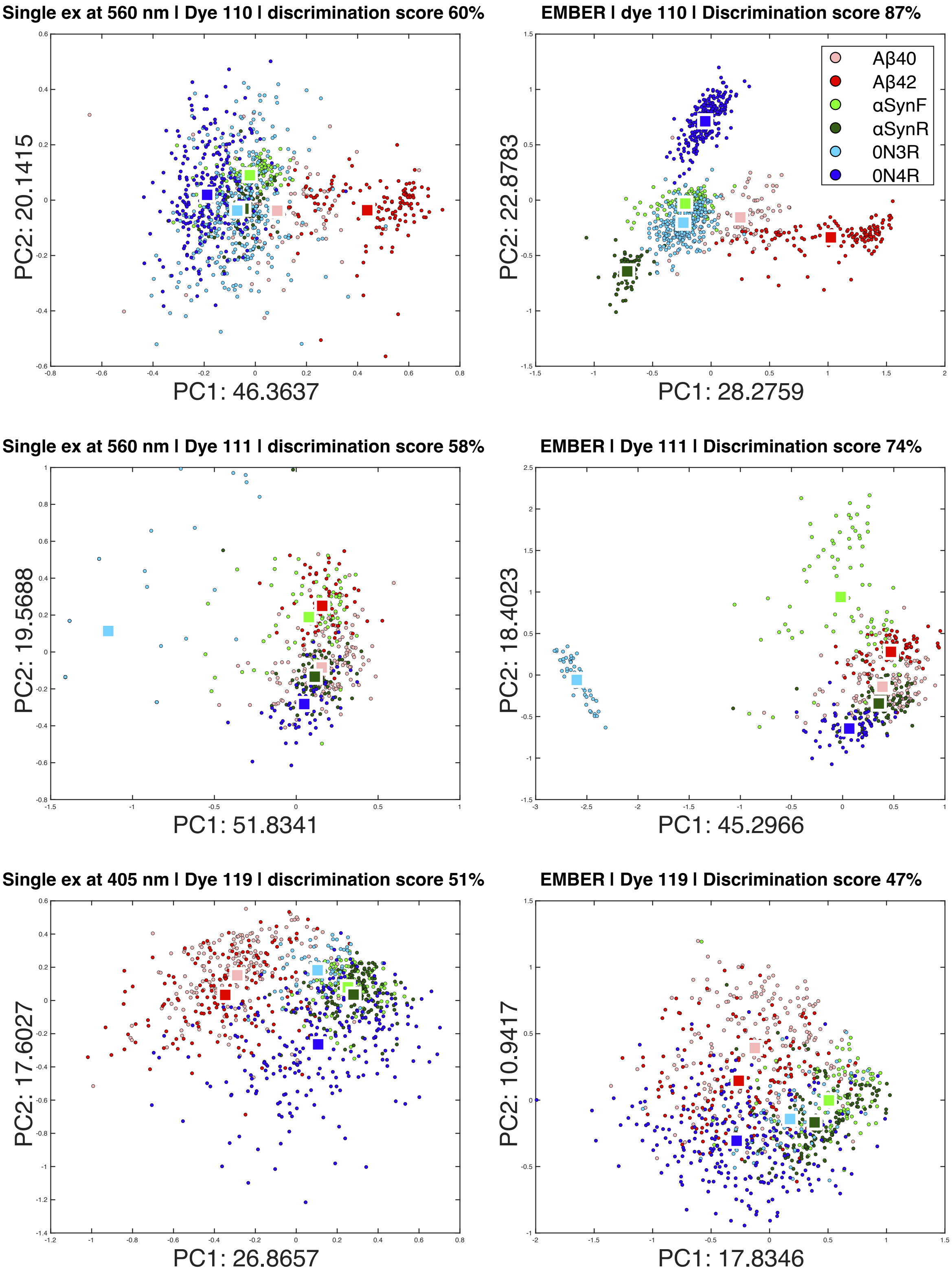


**Fig. S6.** EMBER vs single-wavelength excitation discrimination power comparison for six *in vitro* fibril types. $\lambda_{max}$ single-wavelength excitation data of three dyes (110, 111, 119) from *in vitro* fibril dataset were pulled, and PCA and UMAP analysis were performed (left). The discrimination scores of single-wavelength excitations are lower than that of EMBER (right) representing the outperformance of EMBER in maximizing photophysical property of dyes bound to amyloid fibrils and to discriminate conformational strains of fibrils.

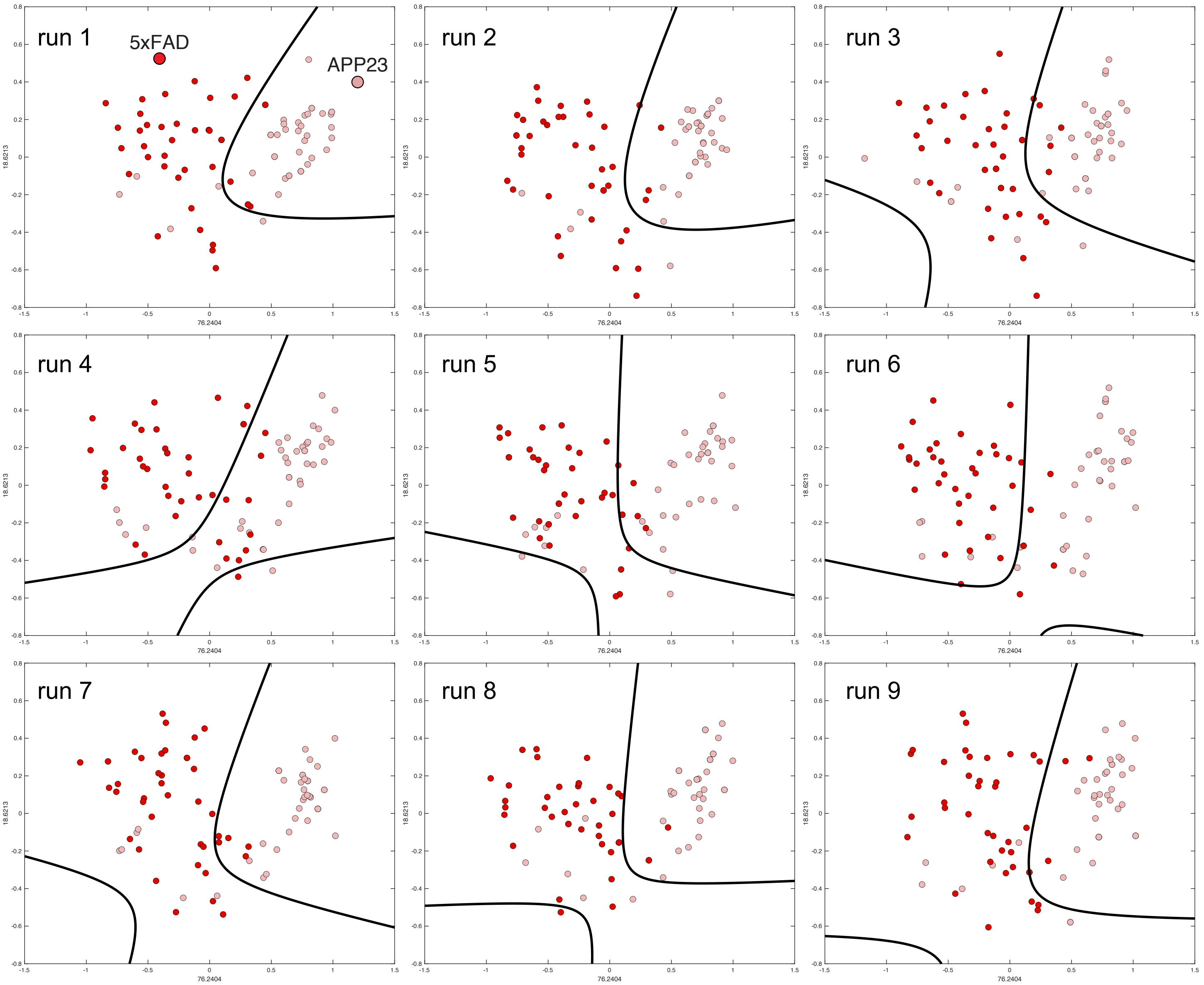


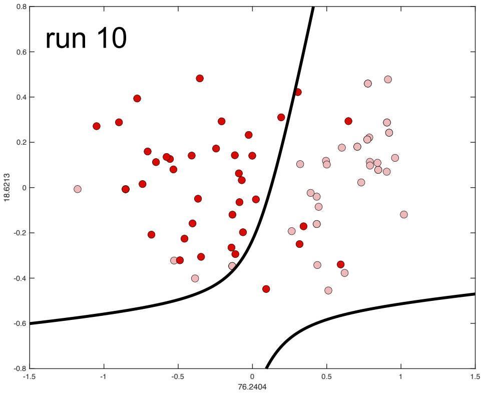


| Run # | Discrimination scores | Average discrimination scores |
|---|---|---|
| run 1 | 89% | 84.3% |
| run 2 | 88% | |
| run 3 | 84% | |
| run 4 | 80% | |
| run 5 | 81% | |
| run 6 | 83% | |
| run 7 | 86% | |
| run 8 | 84% | |
| run 9 | 85% | |
| run 10 | 83% | |

**Fig. S7.** 10 repetitions of quadratic discrimination cluster classification algorithm were performed to quantify discrimination score of plaques in brains of two mouse models. For PCA or UMAP plot, 40 random particles from each fibril sets were concatenated into an array and grouped for quadratic discrimination. Discrimination score is calculated per run, and the average of ten discrimination scores was used as the discrimination score as presented in the main Figure 3. Boundaries pertaining fit discriminants are presented in black lines.

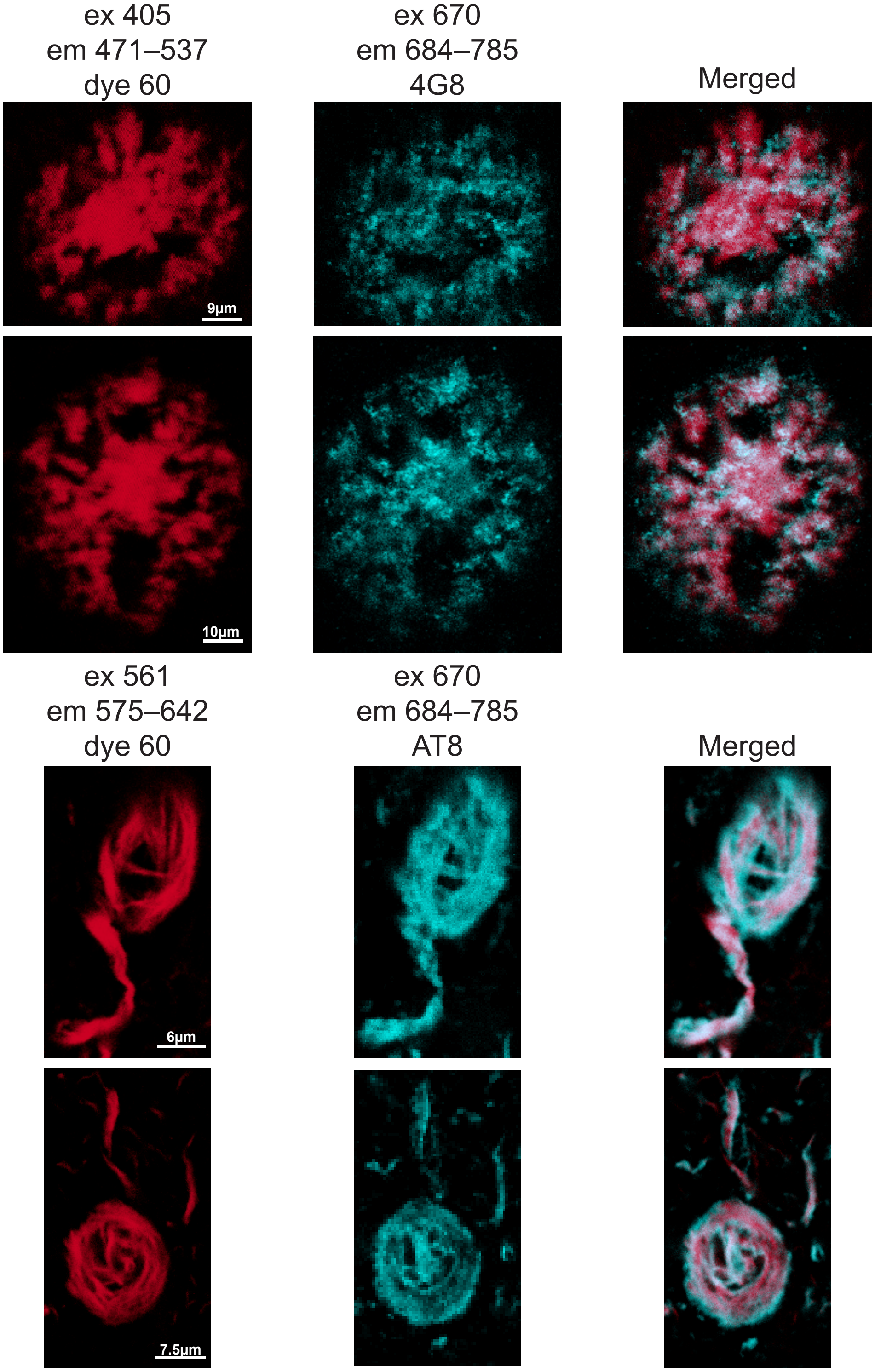


**Fig. S8.** sAD brain donor samples are co-stained with dye 60 and either 4G8 (Aβ) or AT8 (tau) antibody. Dye 60 excites Aβ plaques at 405 nm and tau tangles at 561 nm. The 4G8-stained Aβ plaques or and AT8-stained tau tangles are excited at 670 nm. The merged micrographs show good overlap between dye 60 labeling and corresponding immunostaining. For antibody staining, antigens were first retrieved by autoclaving FFPE sections in 0.01 M Citrate buffer followed by blocking with 10% goat serum. After incubating slides with the primary antibody in 10% goat serum, secondary antibody labelled with Alexa fluor 647 was applied.

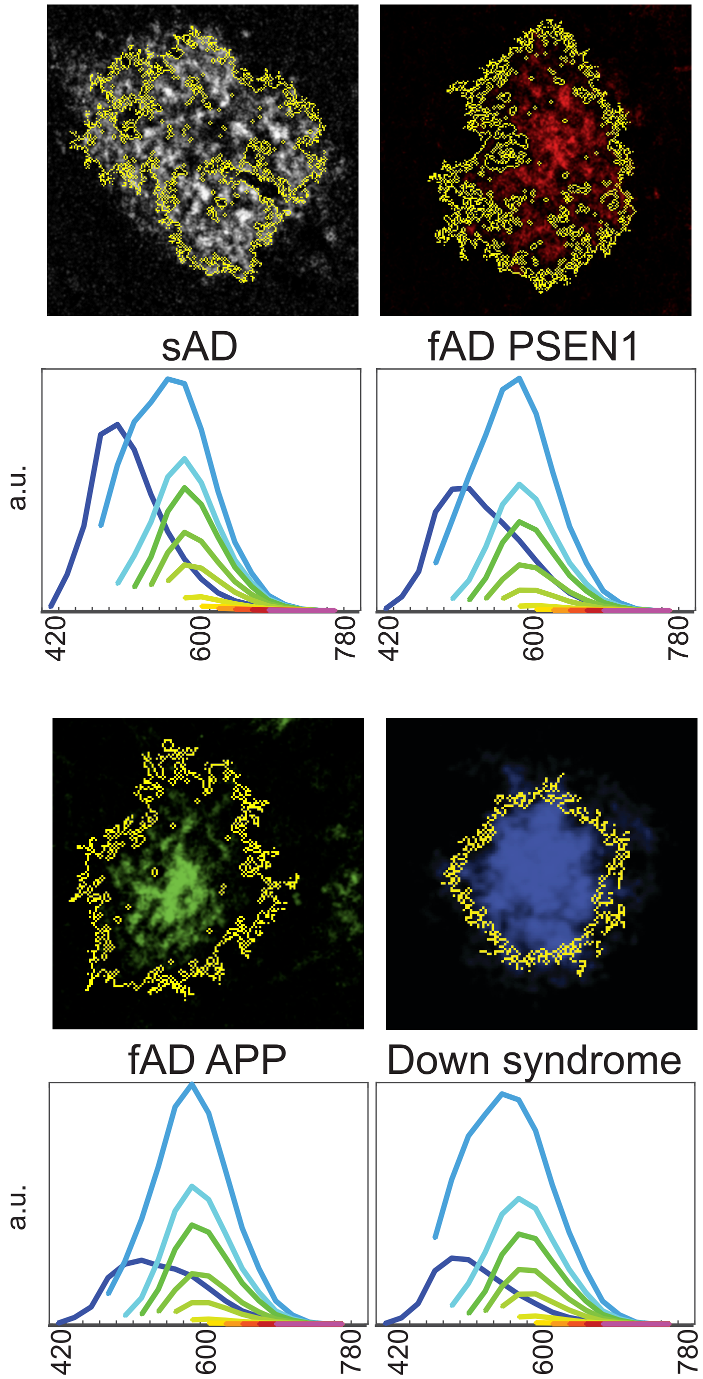


**Fig. S9.** Average EMBER plots for dye 60 stained Aβ plaques across neurogenerative disease brain donor samples. Yellow outlines indicate the particle segmentation performed by customized MATLAB algorithms. Each segmented particle becomes one particle on PCA and UMAP plots.

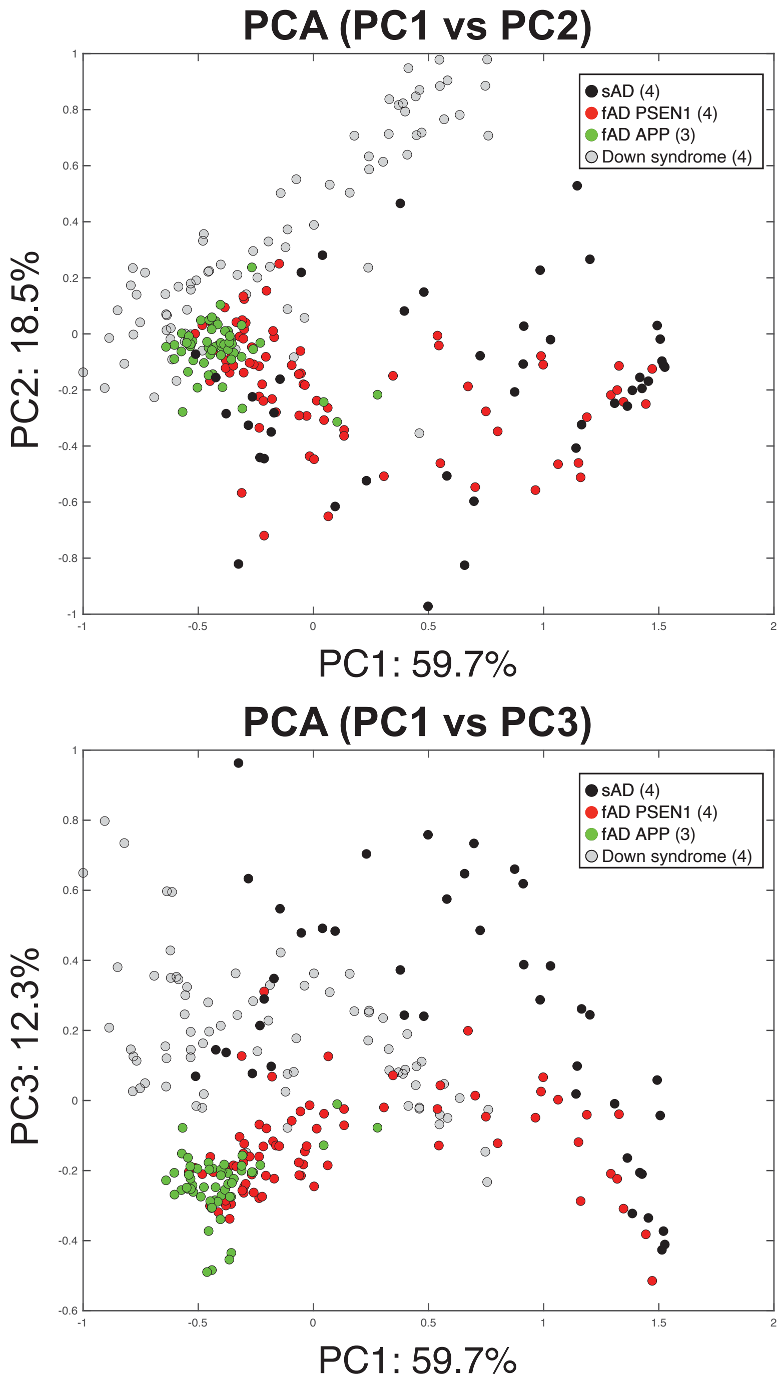


**Fig. S10.** PC1/PC2 and PC1/PC3 PCA plots for Aβ plaques data across neurodegenerative diseases. The number of brain donor samples in each cohort is within the parenthesis.

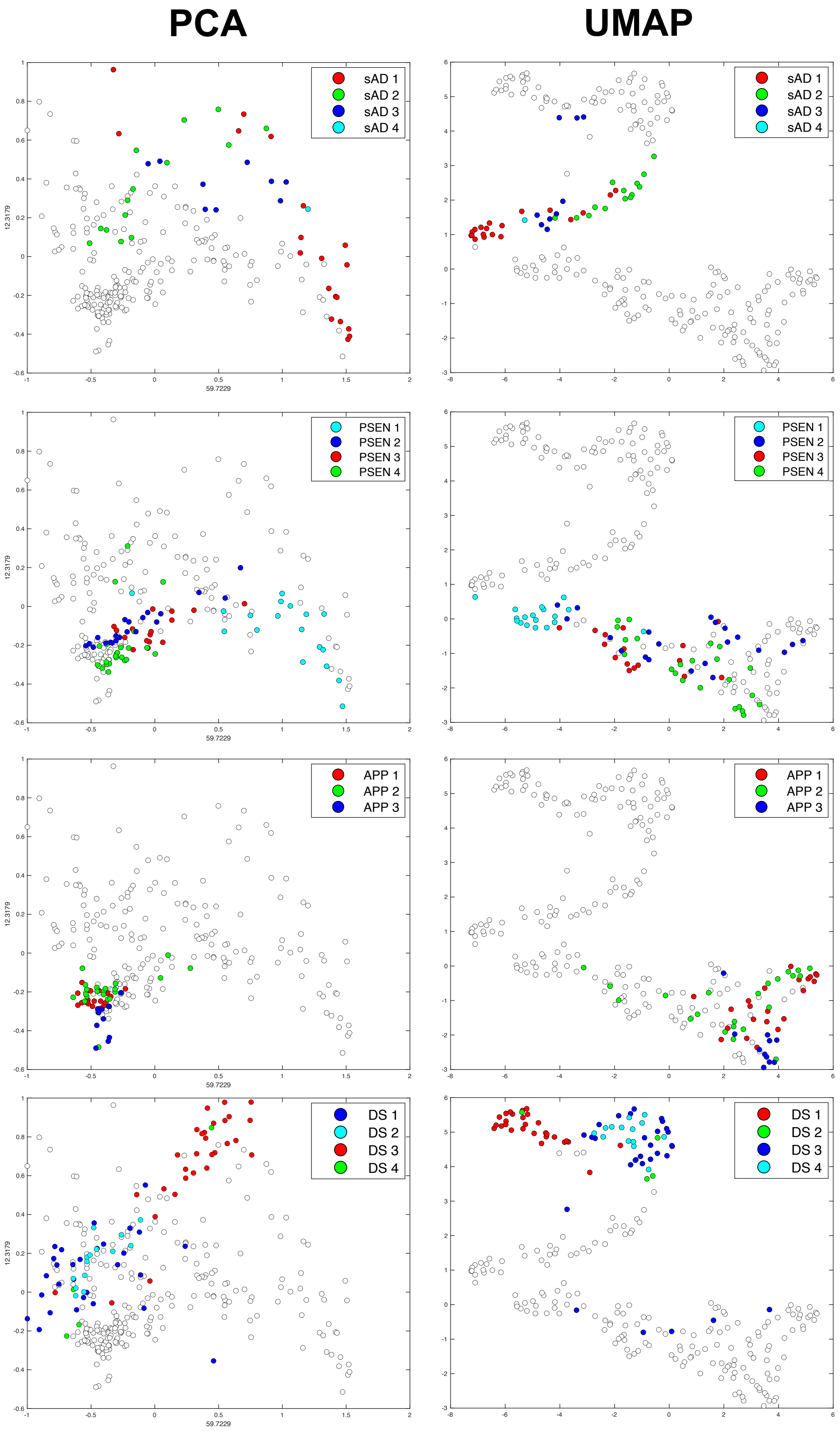


**Fig. S11.** Inter-patient heterogeneity plot for Aβ plaque EMBER data. As shown in main Fig. 6a, Aβ plaques from each patient in each disease cohort were re-plotted using varying colors. To aid visibility, particles pertaining to other EMBER datasets were colored white.

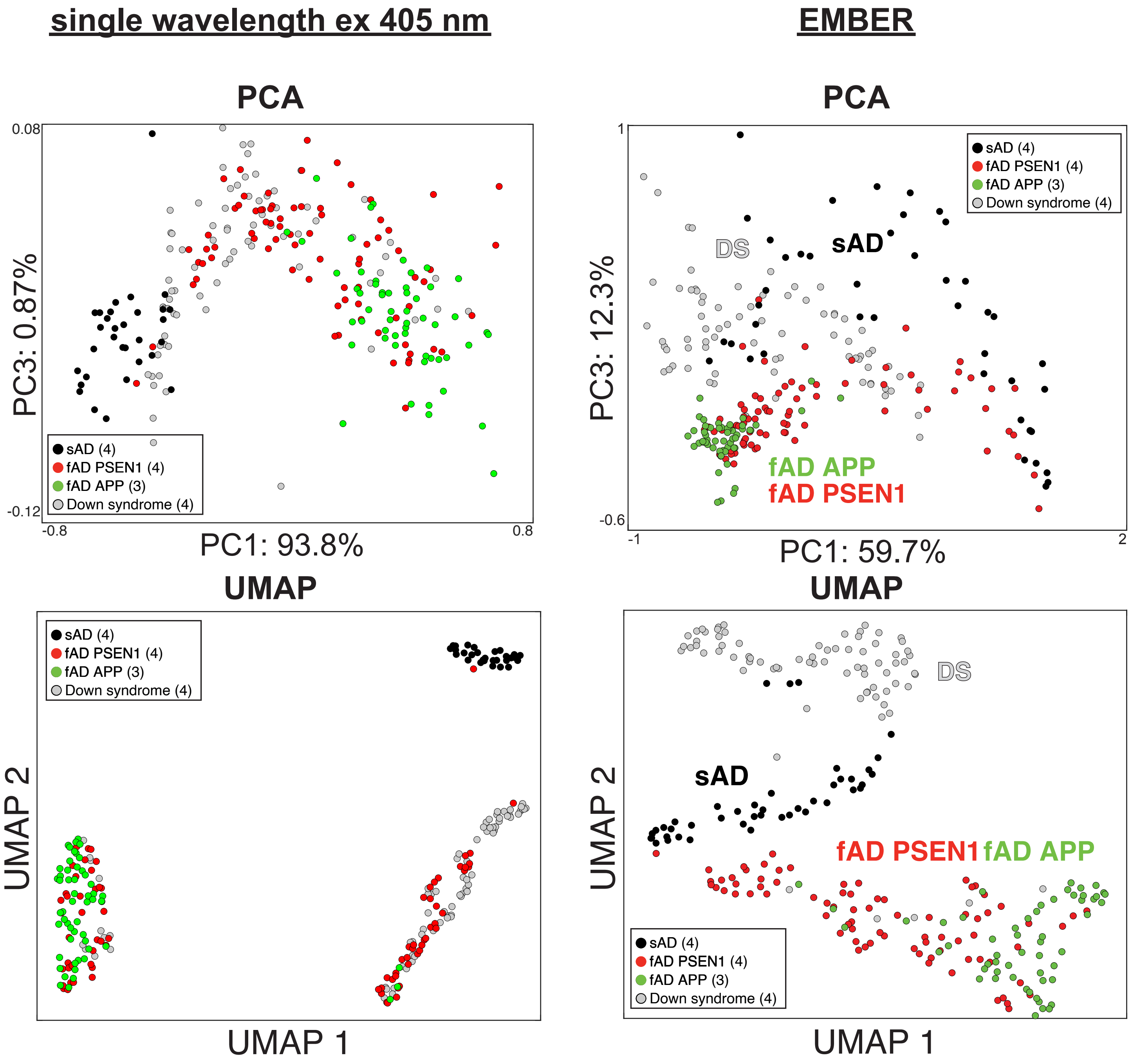


**Fig. S12.** EMBER vs single-wavelength excitation discrimination power comparison for Aβ plaques. 405 nm single-wavelength excitation data was pulled, and PCA and UMAP analysis were performed (left). For single-wavelength excitation, the separation between clusters is worse than that of EMBER (right) representing the outperformance of EMBER in maximizing photophysical property of dyes bound to Aβ plaques and to discriminate conformational strains of fibrils. The number of donor samples in each cohort is within the parenthesis.

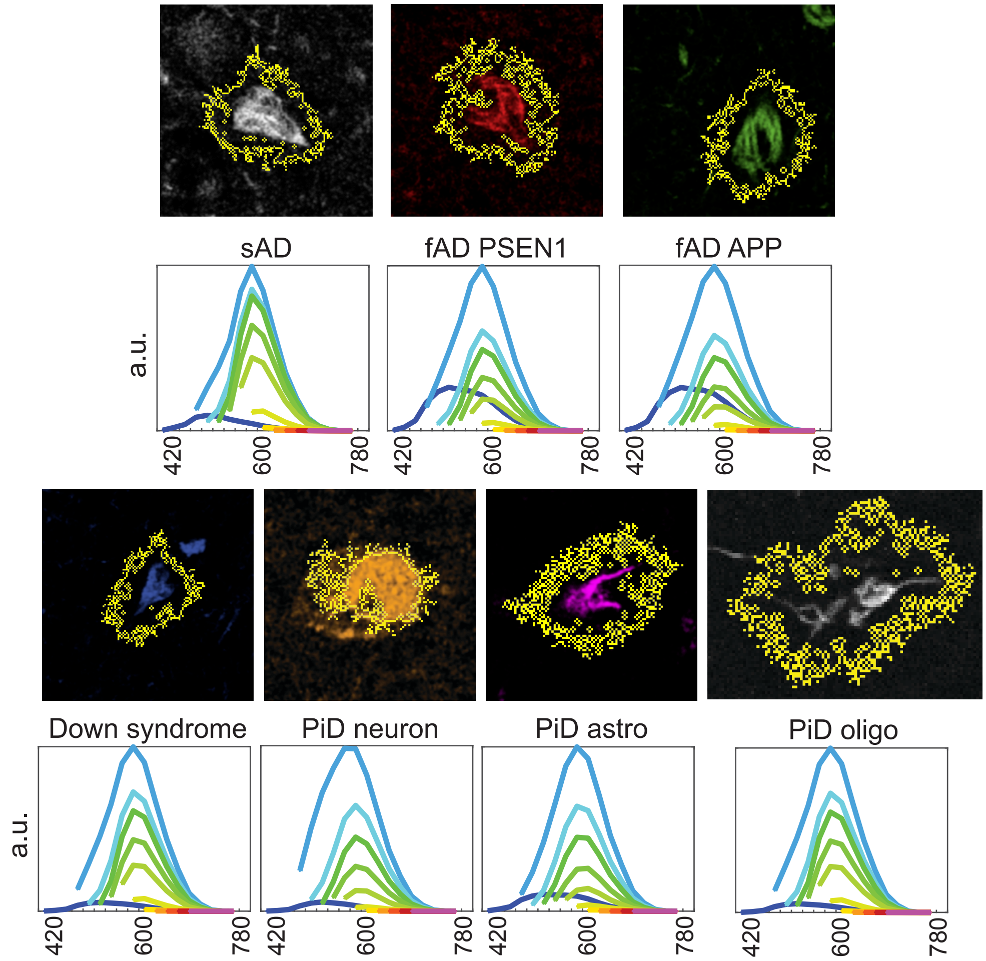


**Fig. S13.** Average EMBER plots for dye 60 stained tau deposits across neurogenerative disease brain donor samples. Yellow outlines indicate the particle segmentation performed by customized MATLAB algorithms. Each segmented particle becomes one particle on PCA and UMAP plots.

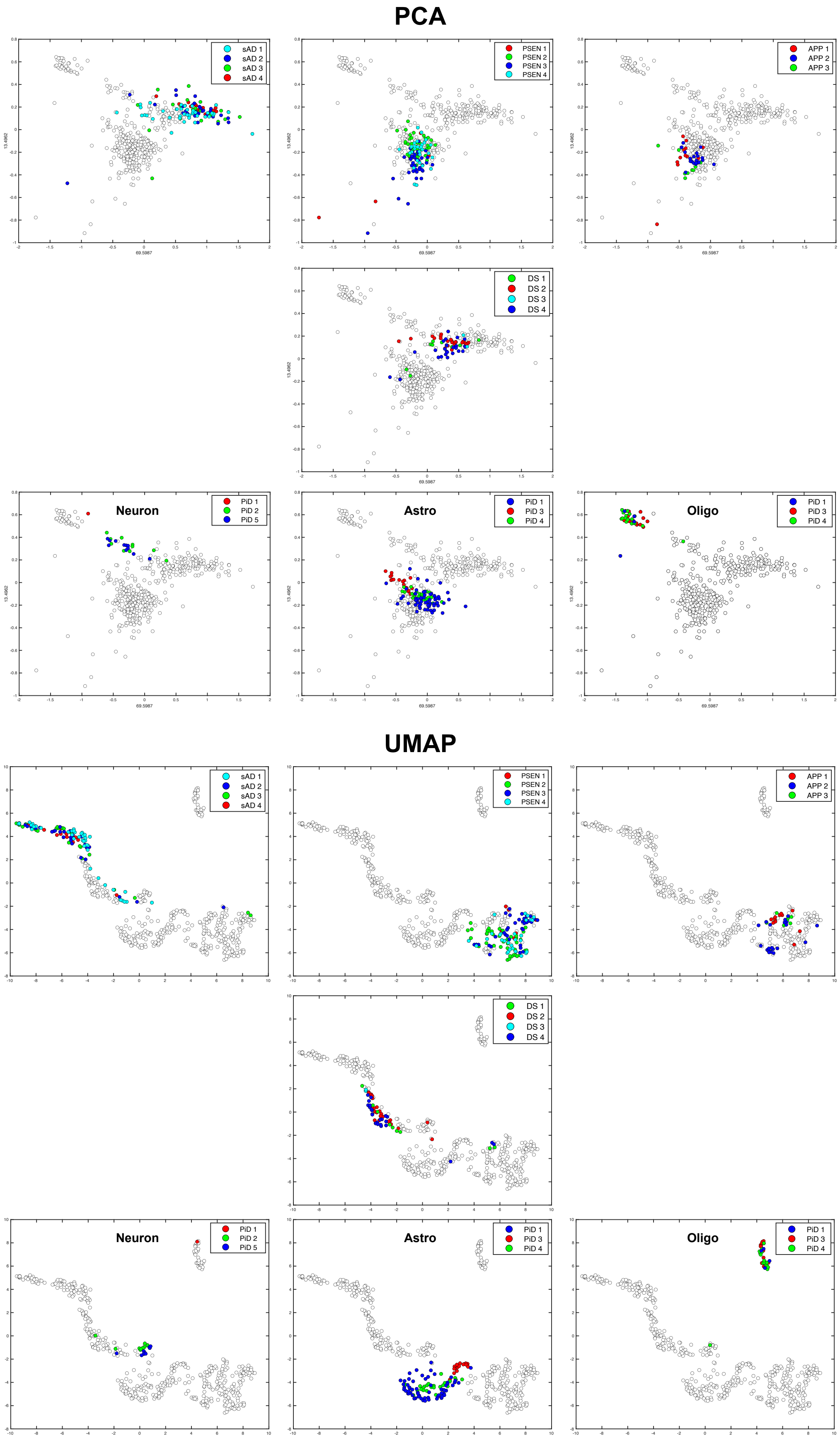


**Fig. S14.** Inter-patient heterogeneity plot for tau deposit EMBER data. As shown in main Fig. 6c, tau deposits of each patient in each disease cohort were re-plotted using varying colors. To aid visibility, particles pertaining to other EMBER datasets were colored white.

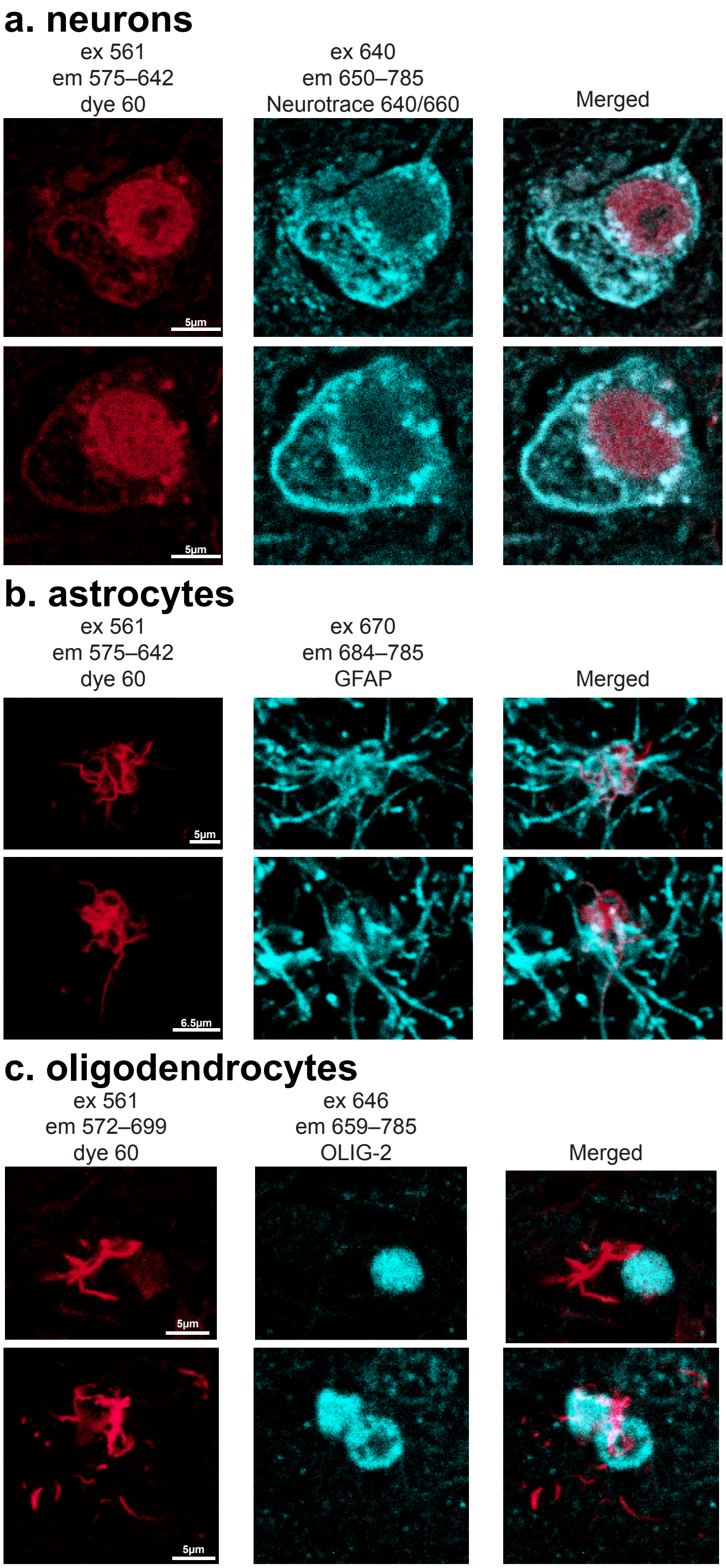


**Fig. S15.** Pick's disease brain donor samples are co-stained with dye 60 and either (a) NeuroTrace640/660 for neuron, (b) GFAP for astrocytes, or (c) OLIG-2 for oligodendrocytes. Dye 60 excites tau tangles of PiD neurons, astrocytes, and oligodendrocytes at 561 nm. Neurotrace640/660 stained neurons excited at 640 nm, GFAP antibody-stained astrocytes excited at 670 nm, and OLIG-2-stained oligodendrocytes at 646 nm. The merged micrographs show good overlap suggesting tau deposit morphology is linked to each cell type. For NeuroTrace640/650, dye 60 and NeuroTrace640/660 were mixed and applied to sample to stain tau tangles and neurons. For GFAP and OLIG-2 antibody staining, antigens were first retrieved by autoclaving slides with 0.01 M Citrate buffer and blocked with 10% goat serum. After incubating slides with primary antibody in 10% goat serum, secondary antibody labelled with Alexa fluor 647 was applied.

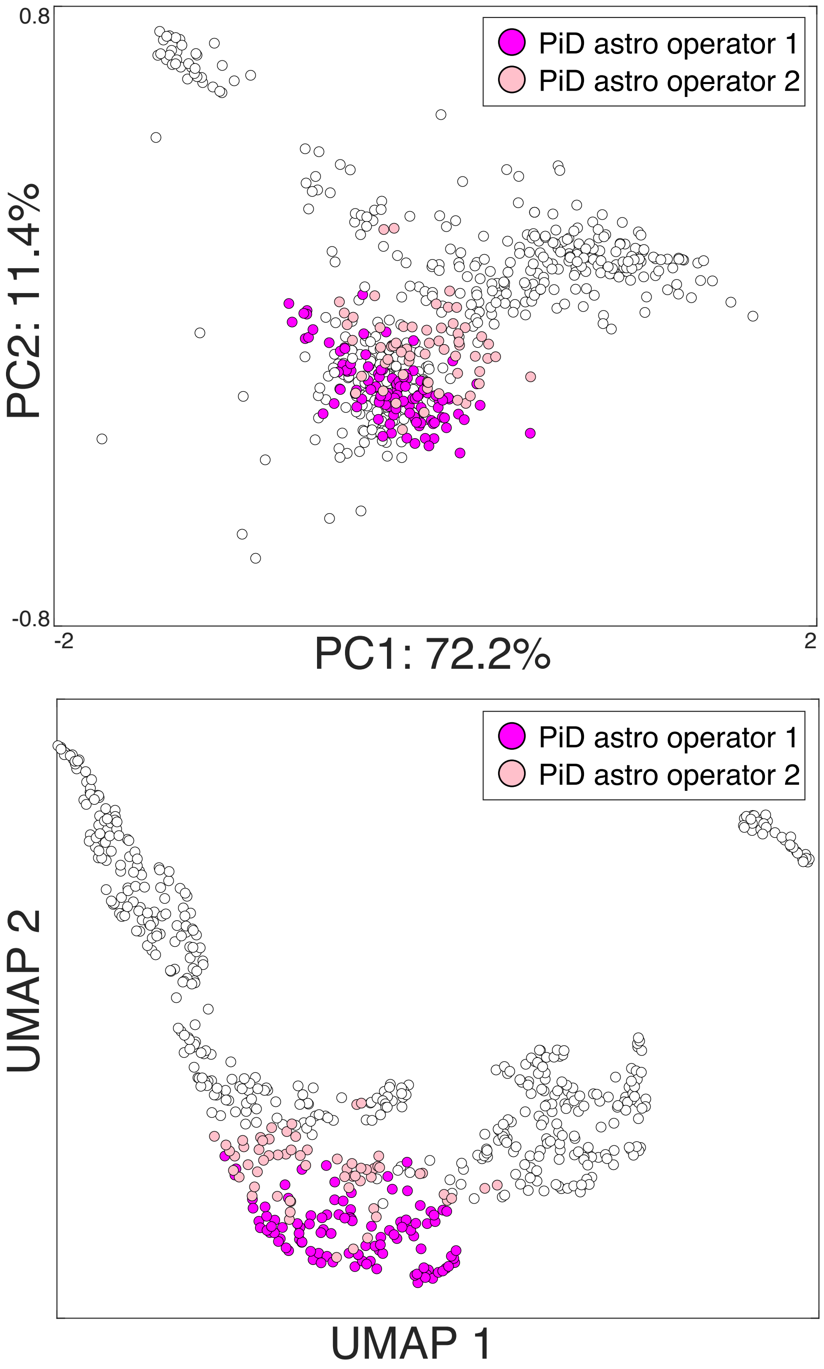


**Fig. S16.** EMBER reproducibility study of PiD tau in astrocytes. Two operators (pink and purple) performed two independent EMBER data collection on three PiD brain donor samples stained with dye 60. The collected data was appended onto the tau deposit dataset as shown in main Figure 6 and PCA and UMAP analysis were performed. The plot shows good overlap between two independent data collections suggesting good reproducibility. To aid visibility of the overlap, the particles pertaining to other EMBER datasets were colored white.

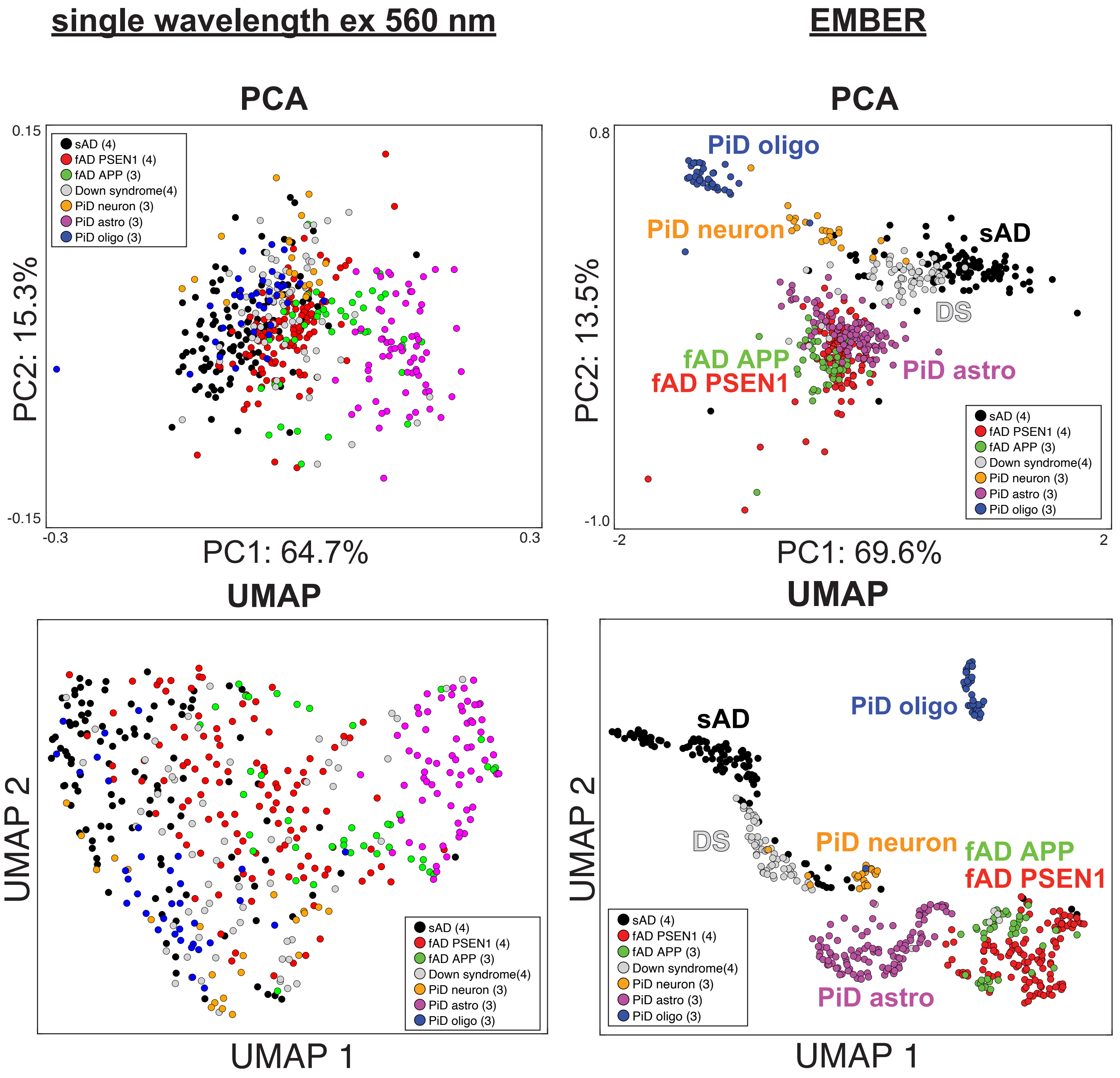


**Fig. S17.** EMBER vs single-wavelength excitation discrimination power comparison for tau deposits. $\lambda_{max}$ single-wavelength excitation data of 560 nm excitation was pulled, and PCA and UMAP analysis were performed (left). For single-wavelength excitation, the separation between clusters is worse than that of EMBER (right) representing the outperformance of EMBER in maximizing photophysical property of dyes bound to tau deposits and to discriminate conformational strains of fibrils. The number of donor samples in each cohort is within the parenthesis.
